# Coevolution of retroviruses with *SERINC*s following whole-genome duplication divergence

**DOI:** 10.1101/2020.02.24.962506

**Authors:** Pavitra Ramdas, Vipin Bhardwaj, Aman Singh, Nagarjun Vijay, Ajit Chande

## Abstract

The *SERINC* gene family comprises of five paralogs in humans of which *SERINC3* and *SERINC5* inhibit HIV-1 infectivity and are counteracted by Nef. The origin of this anti-retroviral activity, its prevalence among the remaining paralogs, and its ability to target retroviruses remain largely unknown. Here we show that despite their early divergence, the anti-retroviral activity is functionally conserved among four human *SERINC* paralogs with *SERINC*2 being an exception. The lack of activity in human *SERINC*2 is associated with its post-whole genome duplication (WGD) divergence, as evidenced by the ability of pre-WGD orthologs from yeast, fly, and a post-WGD-proximate *SERINC*2 from coelacanth to inhibit nef-defective HIV-1. Intriguingly, potent retroviral factors from HIV-1 and MLV are not able to relieve the *SERINC*2-mediated particle infectivity inhibition, indicating that such activity was directed towards other retroviruses that are found in coelacanth (like foamy viruses). However, foamy-derived vectors are intrinsically resistant to the action of *SERINC*2, and we show that a foamy virus envelope confers this resistance. Despite the presence of weak arms-race signatures, the functional reciprocal adaptation among *SERINC*2 and *SERINC*5 and, in response, the emergence of antagonizing ability in foamy virus appears to have resulted from a long-term conflict with the host.

## Introduction

Viruses have been exploiting the host machinery for their persistence, and, in response, the host has continued to evolve increasingly intricate antiviral defense strategies. As part of this ongoing arms-race, while viruses have relied on acquisition and fusion of diverse genes (1), the host-defense mechanism has been made possible by the functional divergence of gene copies following duplication of genes as well as whole-genomes (2–7). Restriction factors being at the forefront of this long-term conflict (8–10), show clear molecular signatures of arms-race (11,12). In fact, the presence of these signatures has been proposed to be a hallmark of restriction factors (8,13), and has been used as a screening strategy to identify potential candidates(2,14). In contrast, the recently identified anti-retroviral host inhibitors *SERINC*5 and *SERINC*3 display an uneventful evolutionary history (15). This is counterintuitive, because distant retroviruses, with wide host-range, encode anti-*SERINC*5 virulent factors (16–18). We, therefore, sought to trace the origins of this antiretroviral activity and its relevance for retroviral inhibition. Our analysis to comprehend the evolutionary origins of the antiretroviral activity in *SERINC*s, identifies an active *SERINC*2 with a hitherto unknown interaction with a foamy virus.

## Results

### Antiretroviral activity among human *SERINC* paralogs

Analysis of sequence similarity and gene structure conservation reveals that *SERINC*5 and *SERINC*4 share a recent ancestry **(Fig-S1)**. Similarly, *SERINC*3 and *SERINC*1 paralogs are most similar to each other (~60% identity). Despite having the lowest sequence similarity to either of the established anti-viral *SERINC* paralogs, *SERINC*2 is relatively similar to *SERINC*3 (50%) and *SERINC*5 (37%). Given such levels of sequence similarity and conserved membrane topology **(Fig-S2)**, we checked the activity of other paralogs to inhibit HIV-1 infectivity in addition to *SERINC*5 and *SERINC*3. From the native versions of *SERINC*1 and *SERINC*4 we failed to express the proteins, so we decided to generate codon-optimized versions of these genes. Having found the proteins being expressed upon codon optimization (Fig1A), we used these conditions to check the inhibitory activity of *SERINC*1 and *SERINC*4. Consistent with previous reports (19,20), we find that *SERINC*2 is the only *SERINC* to have no activity against HIV-1 while all the other *SERINC*s inhibit the virus infectivity to a varying degree (Fig-1A and 1B). Despite less-prominent membrane localization, *SERINC*4 has a potent activity and is second-most powerful in inhibiting the virus (Fig-1C).

**Figure 1.**
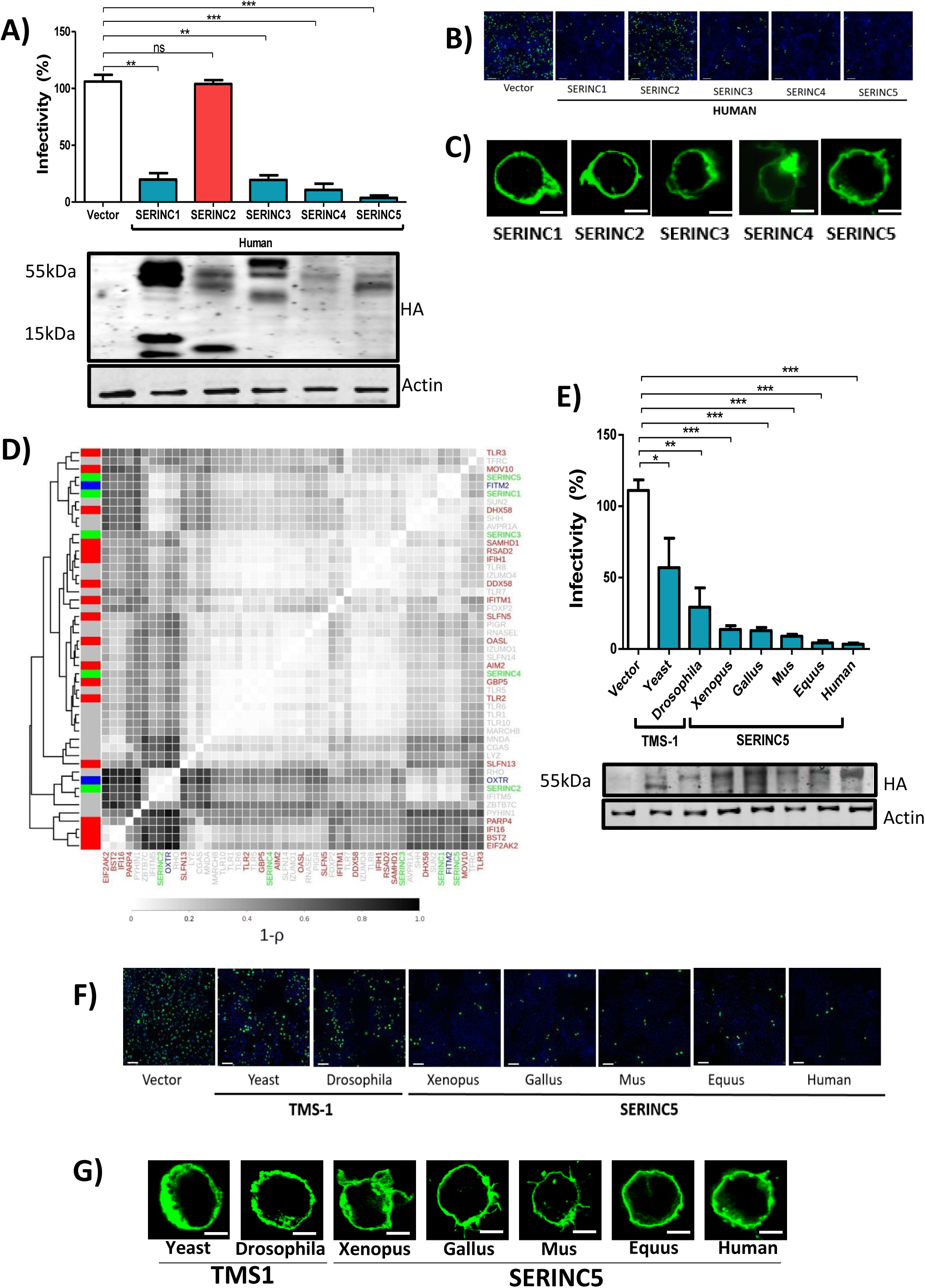
The activity of *SERINC* paralogs and orthologs on HIV-1 infectivity. A. The activity of Human *SERINC* paralogs (*SERINC*1-*SERINC*5) on nef-defective HIV-1 Infectivity. (n=4, mean ± s.d., unpaired t-test, (*)p<0.05 (**)p<0.01 (***)p<0.001, ns-not significant). Lower panel, Western blot showing expression of c-terminally HA-tagged *SERINC* and the corresponding B-actin from cell lysate (HEK293T). Values obtained from the empty vector control was normalized to 100% for comparison with *SERINC* expressors
B. Representative images of TZM-GFP cells infected with viruses produced for Fig-1A. Cells are stained with Hoechst and captured at 10x magnification on CellInsight CX7 High Content Screening platform. Scale-100µm
C. Immunofluorescence assay for the indicated HA-tagged human *SERINC*1-5 transfected in JTAg *SERINC*5/3^-/-^ cells and visualized using Alexa 488 secondary antibody. Scale-10µm
D. Hierarchical clustering of the arms-race signatures in primate genes aligned using ClustalW aligner and used as input for FUBAR. Interferon-induced genes identified by (Shaw et al 2017) are color-coded as red (upregulated), blue(downregulated), green (*SERINC* paralogs), grey (other select genes)
E. The activity of pre-and-post-WGD *SERINC5* orthologs on nef-defective HIV-1 infectivity. (n=4, mean ± s.d., unpaired t-test, (*)p<0.05 (**)p<0.01 (***)p<0.001, ns-not significant). Lower panel, Western blot showing expression of indicated orthologs tagged with HA and the corresponding B-actin from cell lysate (HEK293T)
F. Representative images of TZM-GFP cells infected with viruses produced for Fig-1E. Cells are stained with Hoechst and captured at 10x magnification on Cell Insight CX7 High Content Screening platform. Scale-100µm
G. Immunofluorescence assay for indicated HA-tagged *SERINC* orthologs transfected in JTAg *SERINC*5/3^-/-^ cells and visualized using Alexa 488 secondary antibody. Scale-10µm

In contrast to other restriction factors, *SERINC*5 and *SERINC*3 genes have a rather uneventful history that is distinct from the traditional signatures of recurrent selection seen in genes that are part of an arms-race (21). We investigated the generality of whether the absence of signatures of positive selection was prevalent across all the *SERINC* paralogs. Towards this, we compared the evolutionary signature of *SERINC* genes with previously identified restriction factors, genes showing recurrent positive selection and a few functionally characterized genes that can act as controls (Fig-1D). Within *SERINC* paralogs there is heterogeneity in the patterns of positive selection, with *SERINC*1 and *SERINC*5 showing the least number of positively selected residues while, *SERINC*3 and *SERINC*4 have relatively a greater number of residues under episodic diversifying selection (Fig-1D; **FigS3, S4;** arms-race signature inference is found to be sensitive to the multiple sequence alignment tool used, see **Fig-S5)**. Functionally though, *SERINC*4 and *SERINC*5 are the most potent antiretroviral inhibitors compared to the other three *SERINC*s, suggesting that merely the fraction of sites that are positively selected is not an indicator of its ability to inhibit the retrovirus. Although the arms race signatures of few restriction factors such as BST-2 and EIAF2AK2 were well correlated, the other newly identified restriction factors, including *SERINC*s did not show any consistent pattern of clustering (Fig-1D). One of the prime features of restriction factors has been their ability to get augmented upon interferon stimulation. While the genes which formed a cluster are interferon responsive (IFN responsive genes were obtained from ref (22)), this property did not explain the overall pattern of clustering (Fig-1D). Arms race signatures are not prominent in several interferon-inducible genes and innate immune genes, including TLRs (Fig-1D). Hence the lack of arms race signatures in *SERINC*s is probably not very surprising. The ability to restrict HIV-1 among human *SERINC*1 and *SERINC*4 suggests that this is an evolutionarily conserved feature despite the early divergence of these paralogous copies. Hence, we further decided to investigate the evolutionary origins of this anti-retroviral activity.

### Anti-retroviral activity in an ancient feature among *SERINC*5 orthologs

*SERINC*s plausibly shaped retroviral evolution, as indicated by the parallel emergence of anti-*SERINC*5 activity among diverse retroviral genomes. *SERINC* gene family of restriction factors is conserved across eukaryotic species with evident topological similarity **(Fig-S6)**, so we asked to what extent the ability of *SERINC* orthologs to restrict HIV-1 is conserved. Unicellular eukaryotes and invertebrates have a single copy of the *SERINC* gene (*TMS*1); all these orthologs could restrict *nef*-defective HIV-1 (Fig-1E and 1F), albeit the yeast ortholog having modest activity (2-fold) in comparison to all other orthologs (fold inhibition from 4-36). This lower potency compared to all other orthologs could be attributed to inadequate localization on the membrane or heterologous host expression for yeast and fly *TMS*1 in conditions where the remaining orthologs were adequately expressed on the membrane (Fig-1G). Western blot analysis confirmed the expression of C-terminal HA-tagged TMS-1 and other *SERINC*5 orthologs (Fig-1E, lower panel).

### Coelacanth *SERINC*2 restricts HIV-1

Two rounds of WGD in the ancestor of chordates led to a large increase in the number of genes leading to the acquisition of new functions. This repertoire of genes also provides greater flexibility due to their mutually compensatory functions. Hence post-WGD paralogous copies tend to diversify. A single copy of *SERINC* ortholog (*TMS*1) is present in pre-WGD species and, post-WGD, the number of copies has increased to five in tetrapods and six in bony fishes **(FigS1)**. Our investigation of the pre-WGD ortholog of *SERINC* genes from the yeast (*S. cerevisiae*) and the fly (*D. melanogaster*) *TMS1*, found evidence of anti-retroviral activity (Fig-1E). Functional data on the human paralogs also suggests an ancestrally conserved activity of *SERINC*s in restricting HIV-1, *SERINC*2, however, being an exception (Fig1A). Hence, the human *SERINC2* paralog may have lost the ability to restrict HIV sometime after the WGD (~700 to 400 MYA)(23). It is plausible that the species proximal to the WGD-event might retain a version of *SERINC2* with antiviral activity (Fig-2A). We, therefore, decided to systematically screen *SERINC2* orthologs from post-WGD species at varying levels of sequence divergence from human *SERINC2*. This exercise revealed that Coelacanth *SERINC*2 restricts HIV-1 in conditions where *SERINC*2 isoforms from human did not show any activity (Fig-2B). A shorter isoform of the Human *SERINC*2 (Human-201) was found to be topologically similar (**Fig-S7**) to that of Coelacanth *SERINC*2 but lacked the activity to restrict HIV-1 concurrent with the longer isoform.

**Figure 2.**
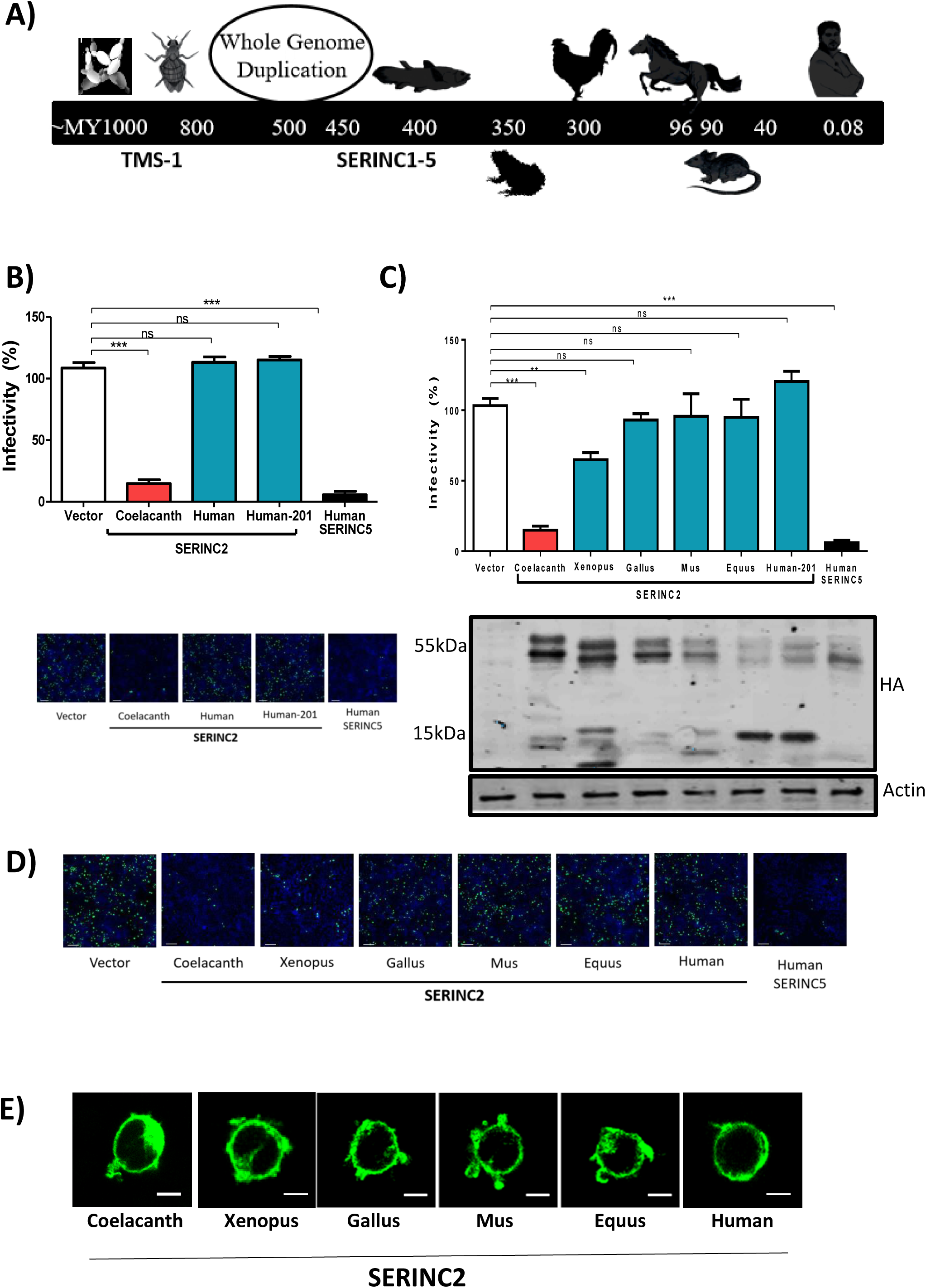
Coelacanth *SERINC*2 inhibits HIV-1 infectivity. A. A Schematic timeline depicting the sequence of events during the course of evolution with species depicted from left to right being *S. cerevisiae, D. melanogaster, L. chalumnae, X. tropicalis, G.gallus, E. caballus, M. musculus* and *H. sapiens*
B. The activity of *SERINC*2 from coelacanth and human *SERINC*2 isoforms on nef-defective HIV-1 infectivity. Values obtained from the empty vector control was normalized to 100% for comparison with *SERINC* expressors. (n=4, mean ± s.d., unpaired t-test, (*) p<0.05 (**)p<0.01 (***)p<0.001, ns-not significant). Lower panel, Representative images of TZM-GFP cells infected with viruses produced for top panel. Cells are stained with Hoechst and captured at 10x magnification on CellInsight CX7 High Content Screening platform.Scale-100µm
C. The activity of *SERINC*2 orthologs from coelacanth, xenopus, gallus, mus, equus and human on nef-defective HIV-1 infectivity. Values obtained from the empty vector control was normalized to 100% for comparison with *SERINC* expressors. (n=4, mean ± s.d., unpaired t-test, (*)p<0.05 (**)p<0.01 (***)p<0.001, ns-not significant). Lower panel, western blot showing expression of *SERINC*2 orthologs tagged with HA and the corresponding B-actin from cell lysate (HEK293T).
D. Representative images of TZM-GFP cells infected with viruses produced for Fig-2C. Cells are stained with Hoechst and captured at 10x magnification on Cell Insight CX7 High Content Screening platform. Scale-100µm
E. Immunofluorescence from for indicated HA-tagged *SERINC* orthologs transfected in JTAg *SERINC*5/3^-/-^ cells and visualized using Alexa 488 secondary antibody. Scale-10µm

### The gradual loss of antiretroviral activity in *SERINC2*

Upon further assessment of anti-HIV-1 activity of post-WGD *SERINC*2 orthologs, we find that while Coelacanth *SERINC*2 reduced the infectivity by ~7-fold, Xenopus *SERINC*2 exhibited a modest inhibition (~2-fold). The activity was completely lost in chicken *SERINC*2 ortholog onwards, this is despite the topology being conserved and the levels of virus-incorporation for chicken ortholog were also appreciable (Fig-2C, **S8 and S9**). This lack of activity, however, is persistent in mouse, horse and human *SERINC*2 (Fig-2C and 2D). Coelacanth *SERINC*2 restricted HIV-1 to a lesser extent compared to human *SERINC*5; however, it was more potent than human *SERINC*3 and *SERINC*1. Further, reduced restriction potency can be attributed to its inadequate membrane presence (Fig-2E), reminiscent of localization of the pre-WGD orthologs from yeast and fly TMS-1 (Fig-1G). We see that there is a dose-dependent inhibition when coelacanth *SERINC*2 was expressed from the plasmids carrying the promoters of different strengths (**Fig-S10**). Western blot of C-terminal HA-tagged *SERINC* expressors verified that each of these *SERINC*2 orthologs is expressed from the synthetic genes (Fig-2C, lower panel). Loss of human *SERINC*2 antiviral activity could have been associated with changes in pathogen repertoire or simply gain of a new function. To experimentally test if this was in response to change in the pathogen repertoire, we asked if the counteraction of Human *SERINC*5 by known retroviral factors, HIV-1 Nef^1^ and MLV GlycoGag^1^, was analogous to that of Coelacanth *SERINC*2. In conditions where Nef and GlycoMA efficiently counteracted the potent restriction imposed by Human *SERINC*5 and the partial restriction of Xenopus *SERINC*2, we find that counteraction of Coelacanth *SERINC*2 restriction by these virulent factors was not appreciable (Fig-3A and 3B). We also inspected if the nef alleles from human and non-human primate lentiviruses showed a similar phenotype and found that nef alleles did not rescue the infectivity comparable to that of *SERINC*5 (Fig-3C). The ability of *SERINC*5 to restrict HIV-1 inhibition varies with the envelope glycoproteins used for pseudotyping. We checked if this is the case with coelacanth SERINC2 as well. Coelacanth *SERINC*2 action indeed phenocopied to that of human *SERINC*5 in terms of the envelope sensitivity (Fig-3D). However, the retroviral factors still failed to rescue this analogous activity of coelacanth *SERINC*2.

**Figure 3.**
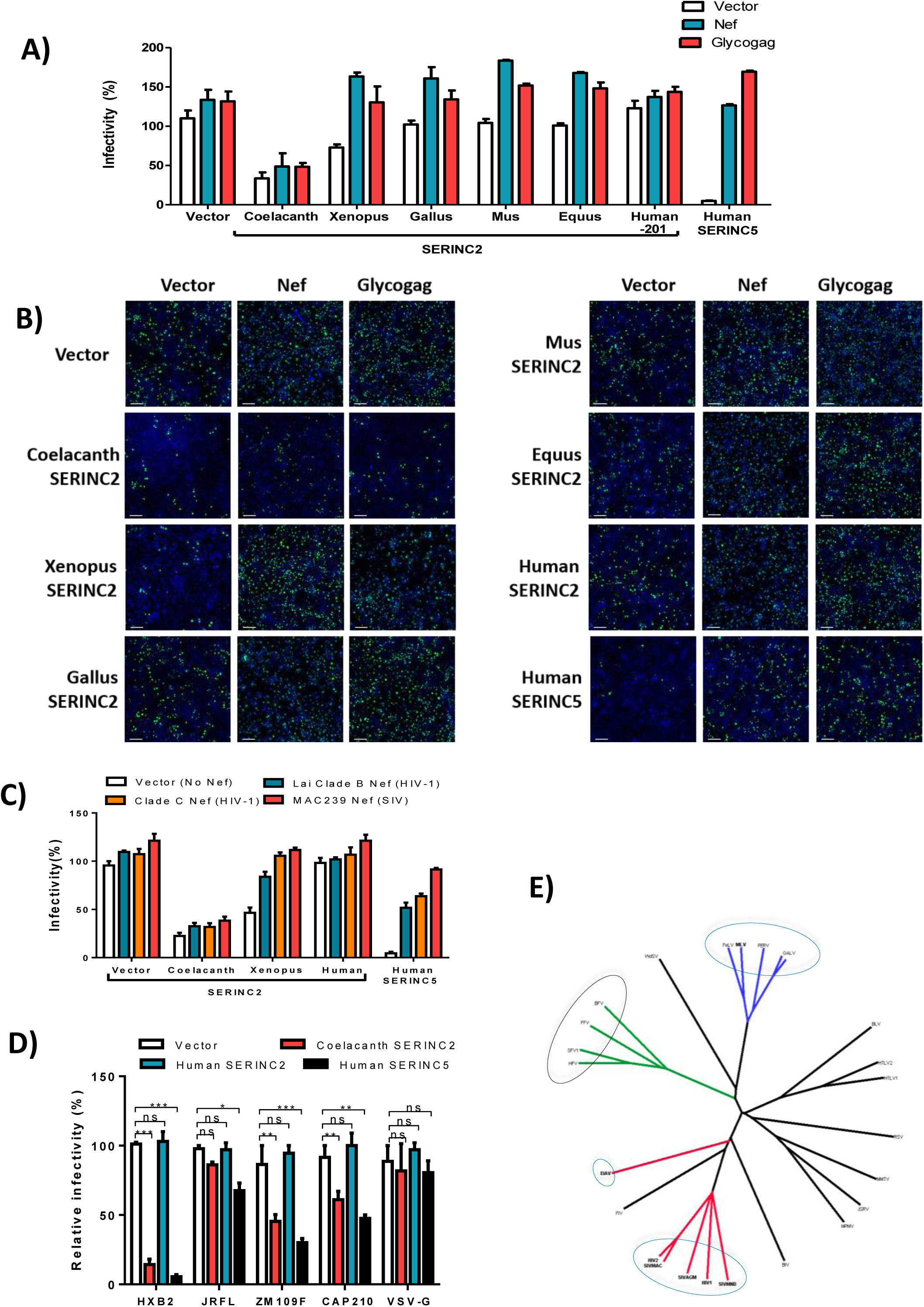
The ability of retroviral factors to antagonize *SERINC*s. A. The ability of Nef and glycoGag to counteract indicated *SERINC*2 orthologs. Human *SERINC*5 was served as control for Nef and glycoGag counteraction of the restriction. Values obtained from the empty vector control without Nef or glycoGag was normalized to 100% for comparison with *SERINC* expressors
B. Representative images of TZM-GFP cells infected for viruses produced for Fig-3A. Cells are stained with Hoechst and captured at 10x magnification on Cell Insight CX7 High Content Screening platform. Scale-100µm
C. The ability of nef alleles to antagonize indicated *SERINC*2 orthologs. Human *SERINC*5 is used as a control
D. Susceptibility of HIV-1 clade-B (HXB2 and JR-FL), clade-C (ZM109F, CAP210) and VSV envelope glycoprotein (VSV-G) to inhibition of infectivity by coelacanth *SERINC*2 with human *SERINC*2 and *SERINC*5 as controls.
E. Phylogenetic tree depicting the relationship among various retroviruses. Unknown retroviral interactions with *SERINC*s are shown in the black ellipse while the known ones are in blue. Branches colored red are complex lentiviruses, blue are gamma retroviruses and green are foamy viruses.

### Functional evidence for arms-race dynamics

Three distinct genera of retroviruses were reported to have independently come up with antagonizing factors to elude the inhibition by *SERINC*5. Having no activity in Nef and Glycogag against coelacanth *SERINC*2 (Fig-3A and 3B) whether represents a new case of functional arms-race dynamics is what we wanted to check next (16–18,24) (Fig-3E). We learnt that coelacanth fish has an endogenous foamy virus (25) the genome organization of which resembled the prototype foamy virus (Fig-4A). The activity of coelacanth *SERINC*2 may have been linked to a foamy virus infection of the host. To experimentally test the idea of whether *SERINC*2 has in-fact evolved to restrict foamy viruses, we checked the presence of anti-foamy virus activity in human *SERINC* paralogs as well as the *SERINC*2 orthologs. Unexpectedly, we find that foamy is insensitive to any of the human *SERINC* paralogs tested as well as the coelacanth *SERINC*2, in conditions where HIV-1 was consistently inhibited (Fig-4B). We argued that this might have been associated with an intrinsic ability of a Foamy virus-encoded factor to counter the inhibition. To dissect this, we co-expressed foamy genes individually to check its ability to rescue, now, *nef*-defective HIV-1 such that the insensitivity of foamy and the presence of antagonizing factor can be revealed. Surprisingly, with none of the foamy components expressed in-*trans* we find an ability to rescue the *nef*-defective HIV-1 in the presence of Human *SERINC*5 (Fig-4C). The inhibition exerted by coelacanth *SERINC*2, however, was antagonized by the Envelope glycoprotein of Foamy virus using an equivalent amount of nef expressing plasmid (Fig-4D), as well as in a dose-dependent manner **(Fig-S11)**. We interrogated if this rescue was an effect of cross-packaging of the HIV-1 core by the foamy-virus envelope glycoprotein. By using a pseudoparticle comprising of an HIV core complemented with the foamy-virus envelope, we show that this was not an effect of cross packaging **(Fig-S12)**, and that this rescue by the envelope glycoprotein was specific to its ability to antagonize coelacanth *SERINC*2 (Fig-4D). To then understand the specificity of this counteraction, we checked the expression of Coelacanth *SERINC*2 at protein level when co-expressed with Foamy envelope. We checked the effect with both low expression (PBJ6) and high expression(pcDNA) of Coelacanth SERINC2 and observed that the steady state levels of Coelacanth SERINC2 is affected in the producer cells when Foamy envelope was co-expressed (Fig-4E). Further, we questioned as to whether the foamy virus was at all sensitive to the effects of coelacanth *SERINC*2. To answer this, we pseudotyped a foamy-virus core, now, with an envelope glycoprotein of HIV-1 lacking a c-terminal domain, for packaging of a heterologous foamy core since the native, full-length, envelope glycoprotein of HIV-1 was not capable of the packaging it. After successful packaging using c-terminally deleted HIV envelope, we show that intrinsically the foamy core is sensitive coelacanth SERINC2 restriction (Fig 4F). This advocates that foamy-virus is indeed sensitive to coelacanth *SERINC*2 and has developed a mechanism to antagonize the inhibitory effect through the envelope glycoprotein likely as part of an ongoing arms-race.

**Figure 4.**
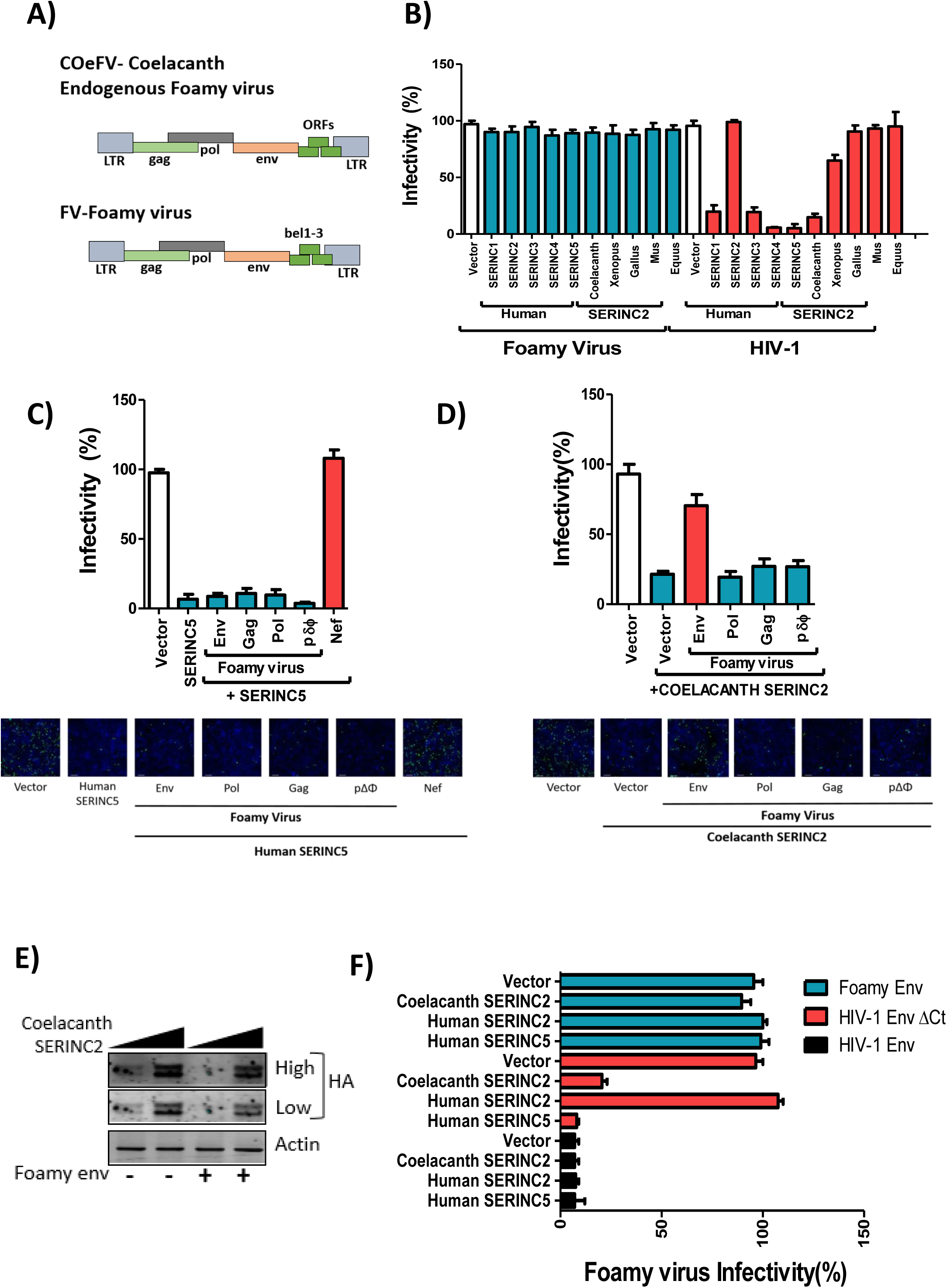
The effect of Coelacanth *SERINC*2 on Foamy virus infectivity and antagonism by the envelop glycoprotein. A. Genome organization of an endogenous foamy virus (CoeFV) from coelacanth and the prototype foamy virus (FV)
B. Infectivity of foamy virus-derived vector and a nef-defective HIV-1 in the presence of indicated human *SERINC* paralogs and *SERINC*2 orthologs. Foamy virus and HIV-1 infectivity obtained from the empty vector control was normalized to 100% for comparison with *SERINC* expressors
C. Nef-defective HIV-1 infectivity obtained from the empty vector control, normalized to 100% for comparison, and with human *SERINC*5. Each foamy virus plasmids expressing different components (gag, pol and env) along with the transfer vector (pδϕ) were used for its ability to counteract human *SERINC*5. SIV MAC239 Nef was used as a control. Lower panel, representative images of TZM-GFP cells infected with viruses produced for Fig-4C. Cells are stained with Hoechst and captured at 10x magnification on CellInsight CX7 High Content Screening platform. Scale-100µm
D. Nef-defective HIV-1 infectivity obtained from the empty vector control, normalized to 100% for comparison, and with Coelacanth *SERINC*2. Each foamy virus plasmids expressing different components (gag, pol and env) along with the transfer vector (pδϕ) were used for its ability to counteract Coelacanth *SERINC*2. Lower panel, representative images of TZM-GFP cells infected with viruses produced for Fig-4D. Cells are stained with Hoechst and captured at 10x magnification on CellInsight CX7 High Content Screening platform. Scale-100µm
E. Western blot showing varying expression of Coelacanth *SERINC*2-HA (from PBJ6 and pcDNA vectors) and the corresponding B-actin from cell lysate (HEK293T) in the presence and absence of Foamy virus envelope expressor.
F. The sensitivity of the foamy core to the action of coelacanth *SERINC*2. The foamy core was trans-complemented with either Foamy envelope, HIV-1 envelope or HIV-1 envelope lacking a c-terminal domain and indicated SERINC was co-expressed to check the ability of individual SERINC to restrict foamy virus. (n = 4, mean ± s.d., unpaired t-test).

## Discussion

The use of comparative evolutionary genetic approaches has continued to enrich our understanding of restriction factor biology for more than a decade(26–30). Reciprocal loss of duplicated genes in different species has been shown to contribute to species-specific differences in susceptibility to pathogens(31–34). Similar to reciprocal gene loss, we find that *SERINC*2 and *SERINC*5 show reciprocal functional adaptation against divergent retroviruses. The ability of these orthologous factors to restrict HIV-1 is remarkable, even in the heterologous hosts (Fig1E, 2C and **Fig-S13**) suggestive of a topology associated feature that remained conserved and contribute to virus restriction. Further, the *SERINC*s from a human can restrict divergent viruses, like MLV(16,17), EIAV(15,18), this only indicates a long-term interaction of these host factors which exerted a strong selective pressure in order to shape retrovirus evolution. While exogenous retroviruses have not yet been reported from pre-WGD species that we considered, the range of targets that a pre-WGD *SERINC* (TMS-1) will have remains to be identified. The restriction on mobility of genetic elements in yeast has earlier been shown conserved in APOBEC(35). While genes such as CCR4 and DHH1 that physically interact with TMS-1 have been implicated in yeast retrotransposon activity (36,37) the mechanism that TMS-1 would manifest on retroelements remains to be investigated. TMS-1 in fly may still be required to regulate the mobility of gypsy retroelements as the gypsy envelope has been shown to pseudotype MoMLV-based vector to efficiently infect fly cells (38,39). Moreover, the analogous mechanism that Rous sarcoma virus uses to produce Pr180gag-pol has been found in yeast Ty-1 transposons (40), makes such existence of retroelement interactions for other host-factors like TMS-1 more probable. Post-WGD, *SERINC* has five copies and this may have been associated with tackling increasing diversity of pathogens during speciation and while doing so retaining its core function. As demonstrated here, with cross-packaging studies we show that *SERINC*2 ortholog, which was thought to be inactive, may potentially have constituted a critical barrier to foamy viruses early on, leading to their divergence over more than 450 MY (25,41). Loss of activity in other *SERINC*2 orthologs may have been associated with neofunctionalization: as suggested by localized sequence divergence **(Fig-S14)**, presence of HNF4alpha binding enhancer **(Fig-S15)** and change in tissue-specific expression patterns **(Fig-S16)** exemplified by expression of *SERINC*2 in the liver of primates. Interestingly, Coelacanth *SERINC*2 inhibited HIV-1 and remained invisible to the most powerful retroviral virulent factors nef (MAC239) and to the glycoGag. Further studies can provide more insights into the lack of activity in these retroviral factors.

The ability of foamy virus envelope to counteract the restriction is first such evidence where a canonical gene product is employed to evade *SERINC* restriction by these atypical retroviruses. Foamy viruses co-diverged with their hosts, and this consistently is also visible in their ability to introduce variations in envelope glycoproteins, among other documented genomic alterations. Such changes perhaps are required for its efficient association with cognate entry receptors or prevent host mechanisms from inhibiting the virus propagation. The nature of many such interactions remains to be clarified due to the unavailability of reagents to study the phenotype in its native context. The counteraction phenotype associated with foamy envelope demonstrated here is not unexpected due to following the reason. It has been earlier suggested that *SERINC*5-mediated particle inhibition varies with the HIV-1 envelope from various isolates (16,42,43); indicating envelope as a determinant for *SERINC*5-sensitivity to particle restriction as also recently demonstrated in (ref 38). Moreover, in the case of EIAV and MLV envelope the ability to resist *SERINC*5 action has been observed (16,18). Constant engagement of *SERINC*5 by the host seems to have therefore acted as a driving force during the evolution of these retroviruses. Having found foamy envelope-dependent activity among coelacanth *SERINC*2 is therefore not surprising and is in agreement with previous reports (44,45) suggesting an ability of envelope to interact with SERINC. As a result, an early challenge by this host factor leading to subsequent divergence of foamy envelope towards insensitivity to *SERINC* restriction. The change in foamy envelope glycoproteins among coelacanth endogenous foamy virus and a prototype foamy attest these facts that there may have been a selective pressure imposed by *SERINC*2. Different SERINC paralogs might have specialized to restrain specific viruses leading to coevolution of viruses in response to specific paralog.

We foresee that weaker signatures, exemplified by *SERINC*s (Fig-1D, **S3**) despite the constant challenge from viruses, could be due to the native functions endowed to such transmembrane proteins where the adaptation through diversification in-response to the pathogen would result in a loss of a core function (46). The poor signatures could also be because the antiretroviral activity is spread out over multiple host genes e.g. as a recent report shows that a Nef-sensitive TIM1 activity is potentiated by *SERINC*5 (47). The host, therefore, can afford such redundancy without having to diversify much. Another example is TLRs where the extracellular domain shows signatures of recurrent positive selection in contrast to the conserved membrane-spanning region (9). Intriguingly, virus specific TLRs are under stronger purifying selection than non-viral TLRs (48) potentially due to the larger number of PAMPs associated with non-viral pathogens. This constraint however may be more pronounced in SERINCs as they are multi-pass transmembrane proteins known to inhibit only retroviruses. This nevertheless is indicative of more *SERINC*-like restriction factors that display poor signatures of arms-race but are functionally active. The core function of *SERINC*s in eukaryotes awaits independent observations nevertheless, the most recent effort in providing structural insights into the fly and human ortholog (44) opens up avenues for structure-inspired biological perturbations.

Despite being conserved, SERINCs have managed to diversify their function against distinct retroviruses. In order to explain these dynamics of *SERINC*-retrovirus evolution, we first collated all the known interactions of SERINCs with retroviral factors (Fig-5A). Since SERINC1 and SERINC4 are not translated from the native transcripts in the conditions that have been probed, we reasoned that their presence might have been associated with extinct retroviral factors. Investigation of SERINC1 and SERINC4 orthologs in diverse species, might be able to decipher the proposed interactions (Fig-5A). In contrast to SERINC1 and SERINC4, we find evidence for SERINC2 being counteracted by retroviral factor from foamy virus. Having investigated all the available data, we decided to propose a model of co-evolution between SERINCs and retroviruses that could explain the diversification of function despite weak signatures of arms-race. Host-driven changes to the viral genome sequence is similar to niche-filling models used to explain evolution of phenotypic traits (49). Hence, we reasoned that divergence of SERINC paralogs following WGD would result in divergence of the niches a virus can occupy. Not surprisingly, we find that viruses have evolved two types of retroviral antagonists to resist SERINC restriction. These types reflect the distinct niches that viruses can occupy. In contrast to previously proposed host-driven models of virus evolution characterized by host adaptation and host-switches, our model suggests the creation of host-niches within the same organism due to divergence of paralogous gene copies. We illustrate our model using the example of post-WGD diversification of interaction between SERINCs and distinct types of retroviral factors (Fig-5B).

**Figure 5.**
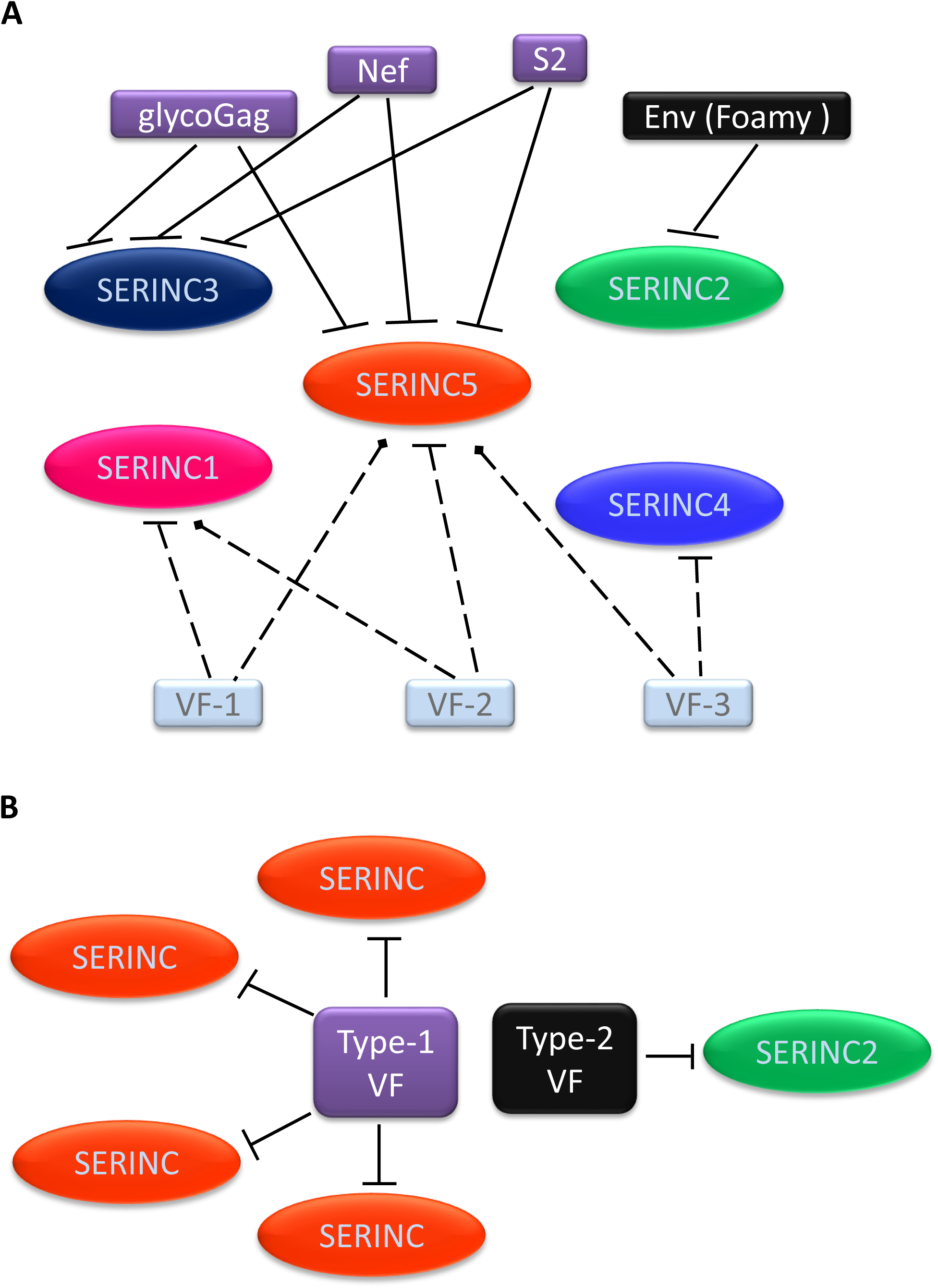
Schematic representation of *SERINC*-retrovirus interactions. A. *SERINC* antagonists (Glycogag, Nef and S2) from distant retroviruses. An interaction between known and hypothesized virulent factors with *SERINC* paralogs. Solid lines show experimentally demonstrated interactions while the dotted lines represent hypothesized interactions. Color-coding for extinct and extant virulent factors (VF). Extinct VFs are shown in light blue, the extant VFs (Type-1) are in purple and Type-2 are in black.
B. A proposed model for adaptive co-evolution among retroviral factors with SERINCs. The evolution of patterns of interactions between viral factors and SERINCs have diversified following post-whole-genome duplication.

In conclusion, evolution-guided analysis for tracking the origin of anti-viral activity in the *SERINC* genes and the dynamics following whole-genome duplication have identified the presence of antiretroviral activity in the only *SERINC* thought to be deficient. The ancient antiretroviral activity among *SERINC*s seems to have challenged retroviruses for more than 450 MY, as suggested by the presence of an evasion strategy in a spumavirus representative to target coelacanth *SERINC*2. The parallel emergence of anti-*SERINC* strategies among divergent retroviruses thus indicates a fundamental role of these host factors in shaping retrovirus evolution.

## Materials and Methods

### Plasmids and reagents

All the plasmids used were isolated using MN NucleoBond® Xtra Midi EF according to the manufacturer’s instructions to minimize endotoxin content. The list of plasmids and reagents used in the experiments are tabulated in ***Supplementary Table I and II***.

### Cell lines

All cell lines used were assessed for mycoplasma contamination and were found to be negative. HEK293T and TZM-GFP indicator cell lines (previously described in Ref (16) were grown in Dulbecco’s Modified Eagle’s Medium (DMEM) with 10% heat-inactivated Fetal Bovine Serum), 2mM L-Glutamine and 1% Penicillin-Streptomycin. Jurkat-TAg^*SERINC*3/5KO^were reported previously (18), maintained in RPMI 1640 with 10% FBS and 2mM L-Glutamine. All the cultures were maintained at 370C and 5% CO2 in a humidified incubator.

### Viruses and infectivity assay

HIV-1 was produced from HEK293T by calcium phosphate transfection and were limited to single-cycle replication. For a 10 cm plate, 7ug NL4-3 env defective and nef defective, 1ug env expressing plasmid and 100ng of pcDNA vectors expressing *SERINC* paralogs and orthologs or equivalent empty vector and 1ug of PBJ6 *SERINC*5 HA was used for virus production. For production of foamy viruses, transient transfection of 0.736 ug pCIES, 1.5ug pCIPS, 11.84ug pCIGS, 11.84 ug pΔΦ and 100ng of pcDNA vectors expressing *SERINC* paralogs and orthologs or equivalent control vector in HEK293T(10 cm plate) was done (50). The viral particles were collected 48 hours post transfection and were centrifuged at 300g for 5 min and quantified by SG-PERT reverse transcription assay (51). Following this, viruses were serially diluted 3, 9 and 27-fold for infection in TZM-GFP reporter cells that were seeded in a 96-well plate, 24 hours prior to infection. HIV-1 Infectivity was quantified by scoring the GFP positive cells using Spectra max MiniMax™ 300 imaging Cytometer (Molecular devices, USA). Foamy Virus infectivity was examined by the number of GFP positive cells indicating the population transduced by Foamy virus. The total cell population was analyzed by nuclei staining with Hoechst 33342 and visualized on CellInsight CX7 High Content Screening platform (Thermo Scientific). The acquired values were normalized with the virus titre obtained from SG-PERT assay as previously described (16) Results are expressed as a percentage of an internal control sample.

### Immunofluorescence

For electroporation in JTag^*SERINC*3/5KO^, cells were grown, and fresh medium was added a day before such that the culture concentration did not exceed 10^6^ cells/ml. Cells in their exponential growth phase were pellet down (10^7^ cells/sample) at 300xg for 10 mins. Prior to adding Opti-MEM, the cells were washed with 1X Phoshphate Buffered Saline (PBS (1X) pH 7.0) to remove residual serum and other cell debris. Each sample was resuspended in 200ul warm OptiMEM. 5ug of constructs expressing HA-tagged *SERINC*2 orthologs or equivalent control vector were then added to the cells. The cells and DNA mixture are added to a 0.2cm gap electroporation cuvette. The cells were pulsed at 140V and 1000uF with exponential decay on BioRAD GenePulser Xcell. 600ul warm RPMI with 20% FBS was immediately added to the electroporated cells, following which they were transferred to a plate containing RPMI with 10% FBS. 48 hours post transfection cells were pellet down at 300g for 5 mins and resuspended in100ul RPMI and laid on poly-L-lysine coated coverslips and fixed with 4% paraformaldehyde. After fixation, the cells were washed twice with 1X PBS. The cells were permeabilized using BD™Perm/wash followed by detection of HA tag with Purified anti-HA.11 Epitope Tag Antibody and Alexa 488 fluorescent tagged secondary antibody. The coverslips were transferred to a glass slide and mounted using ProLong™ Glass Antifade reagent. Images were acquired after 12 hours on Zeiss confocal microscope (LSM740).

### Virus incorporation

Viruses were produced by calcium phosphate transfection in a 10cm plate of HEK293T as mentioned above. The media was replaced with DMEM containing 2% FBS after 12-15 hours of transfection. After 48 hours, the virus containing supernatant was collected and centrifuged at 500xg for 10 mins to exclude any cell fragments. Following this, the viruses were filtered using a 0.22µm syringe filter. This was overlaid on 25% sucrose cushion and concentrated at 50000xg for 2 hours at 4 degrees using a Beckmann Coulter ultracentrifuge. After the spin, the supernatant was aspirated off and the pellet was lysed in Laemmli buffer with TCEP.

### Western blotting

After collection of viruses from producer cells (HEK293T), the cells were collected in 1X ice-cold PBS. They were washed twice at 500g for 5 minutes. The PBS was aspirated until the pellet was completely dry. The pellets were either processed for lysis or stored in −80 until further application. The cell pellets were lysed in DDM Lysis buffer (100mM NaCl, 10mM HEPES pH 7.5, 50mM Tris(2-carboxyethyl) phosphine hydrochloride (TCEP), 1% n-Dodecyl-β-D-maltoside (DDM), 2xcOmplete™, EDTA-free Protease inhibitor cocktail and rocked on ice for 30 minutes. Following this, the lysates were centrifuged at 10000g for 15 mins and the supernatant was collected and mixed with 4x Laemmli buffer with 10mM TCEP.

SDS-PAGE was used for resolving cell pellets and virions for analysis by western blot. The proteins from the gel were transferred onto a PVDF membrane on a semi-dry transfer unit (Hoefer TE77) for 75 minutes with 125 Amp constant current. The membrane containing the proteins was then blocked with 2x Odyssey Blocking Buffer that was diluted with 1x TBS to make 1x Blocking buffer. After 20 minutes of blocking, Anti-HA.11 Epitope Tag Antibody (mouse) and anti-actin(rabbit) (diluted 1: 1000 in 1x blocking buffer) was used to probe the membrane to detect the *SERINC* orthologs and paralogs that were tagged with C-terminal HA tag. After incubation with the primary antibody for 1 hour, the membrane was washed thrice with 1X TBST for 5 mins. Goat anti-mouse 680 and goat anti-rabbit 800 Li-Cor antibodies were used to detect infra-red dyes signal from the membrane. The western blot was analysed on Odyssey imager system.

### Reconstruction of coelacanth *SERINC*5 sequence

The *SERINC*5 from coelacanth was found to be shorter than its annotation from its orthologs. Upon closer inspection it was found that the exons orthologous to human exons1-3 were missing in the coelacanth *SERINC*5 genomic sequence. We found that the upstream region of coelacanth *SERINC*5 had a gap in the genomic sequence. It was also found that the transcribed regions upstream of the gap did not correspond to *SERINC*5 exons. Hence, we decided to use a chromosome walking strategy to reconstruct the missing exons, such that at least at sequence level the topology of the protein could be studied (suppl material for reconstruction of *SERINC*5).

### Data and code availability

The multiple sequence alignments are available for download under GNU license from the github repository https://github.com/ceglab/RestrictionFactorsArmsRace.

### Genome correction and quality check

In order to rule out the possibility of artefacts in evolutionary analysis that arise from errors in the sequence of genome assemblies (true even for high quality reference genomes found on ensemble(52)) we systematically assessed the quality of each of the primate genomes used in this study. Genome assemblies of 15 primate species were downloaded from ensembl release 98 through the ftp site. Whole genome sequencing datasets corresponding to each of these species were obtained from the Short Read Archive (SRA) with the criteria that they should have >~30x coverage. Details of the genome assemblies used and the corresponding raw read data from each species are provided in GitHub repository. The raw read data were mapped to the corresponding genomes using the bwa mem read mapper(53) with default settings. The alignment files obtained from the mapping step were used to generate the genotype likelihood estimates using the program angsd (54). The genotype likelihood estimates were provided to the program referee(55) to assign quality scores and perform genome correction. Overall, we found that in all the primate genomes considered, less than 1% of the bases were corrected by the program referee. The sequencing data used for performing genome correction are not from the same individual that was used for genome assembly. Hence, it is possible that many of the corrected positions are merely nucleotide polymorphisms.

### Manual curation and multiple sequence alignment

The manually curated open reading frame multi-fasta files consisting of 70 genes from ~15 primate species were collected from Ensembl. We extend our previous multiple sequence alignment strategy(56) by including additional multiple sequence alignment programs to generate 8 independent alignments for each gene. Several multiple sequence alignment tools were used to ensure that the inferred patterns of sequence evolution are not restricted to the alignment strategy used. The choice of the actual multiple sequence alignment tools used was based on the performance-based classification of algorithms (57).

### Use of FUBAR to find evolutionary fingerprints

Traditional approaches that endeavour to find arms race signatures in genes look for recurrent occurrence of positive selection in the same gene in different evolutionary lineages. A more recent approach has been to use the full joint distribution of synonymous (α) and nonsynonymous (β) rates as an evolutionary fingerprint of a gene(21). The program FUBAR(58) is available as part of the HyPhy package(59). Pairs of conditionally dependent sites were identified using the program BGM(60). Estimates of selection coefficients are inferred from the multiple sequence alignments for pairs of α-β combinations that form a discretized grid. We calculated the distance measure defined by Murrel(21) to perform hierarchical clustering of the selection signatures obtained from FUBAR.

## Acknowledgements

This work is supported by IYBA fellowship (BT/010/IYBA/2017/01), a grant (BT/PR26013/GET/119/191/2017) from the Department of Biotechnology, Government of India, and the Wellcome Trust/DBT India Alliance Fellowship [grant number IA/I/18/2/504006 awarded to AC]. PR, VB and AS are supported by a fellowship from the MHRD, CSIR and DBT respectively. The computational analyses were performed on the Har Gobind Khorana Computational Biology cluster. Authors thank Jeremy Luban, and the NIH AIDS Reagent Program for the reagents and cell lines. Authors are grateful to Massimo Pizzato for his critical inputs and reagent support.

## Supplementary Figure Legends

**Figure S1-** Phylogenetic analysis of SERINC paralogs. *S. cerevisiae, C. elegans, and D. melanogaster* has a single copy of this gene. Whole-genome duplication that occurred in an early ancestor of mammals gave rise to a cluster of five genes divided into clusters of SERINC1, SERINC2, SERINC3, and SERINC4 and SERINC5. Colors denote different branches of each SERINC in chordates. The lengths of the triangles are proportional to the number of nucleotide substitutions that have taken place in a particular branch. The tree was generated using Ensembl.

**Figure S2-** Topology of human SERINC Paralogs predicted using TOPCONS

**Figure S3**- Arms-race signatures of primate SERINC genes inferred using FUBAR

**Figure S4**- Distribution of sites under different selection regimes across the SERINC genes. The colors are inner (Blue), outer (Pink), helix (Red). Black dots are estimates of dS and red dots are dN. Pairs of conditionally dependent sites identified by the program BGM (Bayesian Graphical Model) are connected by a dotted line (see SERINC1 and SERINC4). Vertical green lines correspond to the sites under purifying selection.

**Figure S5**- Effect of different aligners on hierarchical clustering of arms-race signatures of primate genes. Interferon induced genes identified by (Shaw et al 2017) are color coded as red (upregulated), blue(downregulated), green (SERINC paralogs), grey (other select genes)

**Figure S6**- Topology of SERINC5 orthologs predicted using Topcons. Coelacanth SERINC5 was reconstructed from the RNAseq data (*) see supplemental material for details

**Figure S7**- Topological features of Human SERINC2 splice isoforms and coelacanth SERINC2 predicted using TOPCONS

**Figure S8**- Topology of SERINC2 orthologs predicted using TOPCONS

**Figure S9**- Incorporation of SERINC2 orthologs in the virus particles and detection by indicated antibodies against target proteins

**Figure S10**- Effect of a dose-dependent expression of coelacanth SERINC2 on HIV-1 infectivity

**Figure S11**- Counteraction of coelacanth SERINC2 restriction by a Foamy virus envelope glycoprotein

**Figure S12**- Reciprocal-packaging of a foamy envelope with an HIV core and sensitivity to SERINCs

**Figure S13**- Infectivity of HIV-1 produced from JTAgSERINC5/3KO having ectopic expression of the indicated SERINC

**Figure S14**- Multiple sequence alignment of SERINC2 orthologs; highlighted regions showing the sites of sequence divergence

**Figure S15**- Evidence for the presence of HNF4a binding site in the intron of human SERINC2 gene

**Figure S16**- Gene expression evolution of SERINC genes in vertebrates

## References

1. Koonin E V., Dolja V V., Krupovic M. Origins and evolution of viruses of eukaryotes: The ultimate modularity. Virology. 2015.

2. Malfavon-Borja R, Sawyer SL, Wu LI, Emerman M, Malik HS. An evolutionary screen highlights canonical and noncanonical candidate antiviral genes within the primate TRIM gene family. Genome Biol Evol. 2013;

3. Paparisto E, Woods MW, Coleman MD, Moghadasi SA, Kochar DS, Tom SK, et al. Evolution-Guided Structural and Functional Analyses of the HERC Family Reveal an Ancient Marine Origin and Determinants of Antiviral Activity. J Virol. 2018;

4. Daugherty MD, Schaller AM, Geballe AP, Malik HS. Evolution-guided functional analyses reveal diverse antiviral specificities encoded by ifit1 genes in mammals. Elife. 2016;

5. Sackton TB, Lazzaro BP, Clark AG, Wittkopp P. Rapid expansion of immune-related gene families in the house fly, musca domestica. Mol Biol Evol. 2017;

6. Kasahara M. Genome Duplication and T Cell Immunity. In: Progress in molecular biology and translational science. 2010. p. 7–36.

7. Dehal P, Boore JL. Two Rounds of Whole Genome Duplication in the Ancestral Vertebrate. Holland P, editor. PLoS Biol. 2005 Sep;3(10):e314.

8. Duggal NK, Emerman M. Evolutionary conflicts between viruses and restriction factors shape immunity. Nat Rev Immunol. 2012;12(10):687–95.

9. Wlasiuk G, Khan S, Switzer WM, Nachman MW. A history of recurrent positive selection at the toll-like receptor 5 in primates. Mol Biol Evol. 2009;

10. Daugherty MD, Young JM, Kerns JA, Malik HS. Rapid Evolution of PARP Genes Suggests a Broad Role for ADP-Ribosylation in Host-Virus Conflicts. PLoS Genet. 2014;

11. Boso G, Buckler-White A, Kozak CA. Ancient Evolutionary Origin and Positive Selection of the Retroviral Restriction Factor Fv1 in Muroid Rodents. J Virol. 2018;

12. Compton AA, Malik HS, Emerman M. Host gene evolution traces the evolutionary history of ancient primate lentiviruses. Philosophical Transactions of the Royal Society B: Biological Sciences. 2013.

13. Harris RS, Hultquist JF, Evans DT. The restriction factors of human immunodeficiency virus. Journal of Biological Chemistry. 2012.

14. McLaren PJ, Gawanbacht A, Pyndiah N, Krapp C, Hotter D, Kluge SF, et al. Identification of potential HIV restriction factors by combining evolutionary genomic signatures with functional analyses. Retrovirology. 2015;

15. Firrito C, Bertelli C, Vanzo T, Chande A, Pizzato M. SERINC5 as a New Restriction Factor for Human Immunodeficiency Virus and Murine Leukemia Virus. Annu Rev Virol. 2018;

16. Rosa A, Chande A, Ziglio S, De Sanctis V, Bertorelli R, Goh SL, et al. HIV-1 Nef promotes infection by excluding SERINC5 from virion incorporation. Nature. 2015;526(7572):212–7.

17. Usami Y, Wu Y, Göttlinger HG. SERINC3 and SERINC5 restrict HIV-1 infectivity and are counteracted by Nef. Nature. 2015;526(7572):218–23.

18. Chande A, Cuccurullo EC, Rosa A, Ziglio S, Carpenter S, Pizzato M. S2 from equine infectious anemia virus is an infectivity factor which counteracts the retroviral inhibitors SERINC5 and SERINC3. Proc Natl A cad Sci U S A. 2016;113(46).

19. Schulte B, Selyutina A, Opp S, Herschhorn A, Sodroski JG, Pizzato M, et al. Localization to detergent-resistant membranes and HIV-1 core entry inhibition correlate with HIV-1 restriction by SERINC5. Virology. 2018;

20. Sood C, Marin M, Chande A, Pizzato M, Melikyan GB. SERINC5 protein inhibits HIV-1 fusion pore formation by promoting functional inactivation of envelope glycoproteins. J Biol Chem. 2017;292(14).

21. Murrell B, Vollbrecht T, Guatelli J, Wertheim JO. The Evolutionary Histories of Antiretroviral Proteins SERINC3 and SERINC5 Do Not Support an Evolutionary Arms Race in Primates. J Virol. 2016 Jun 29;

22. Shaw AE, Hughes J, Gu Q, Behdenna A, Singer JB, Dennis T, et al. Fundamental properties of the mammalian innate immune system revealed by multispecies comparison of type I interferon responses. PLoS Biol. 2017;

23. Lynch M. The Age and Relationships of the Major Animal Phyla. Evolution (N Y). 1999;

24. Ahmad I, Li S, Li R, Chai Q, Zhang L, Wang B, et al. The retroviral accessory proteins S2, Nef, and glycoMA use similar mechanisms for antagonizing the host restriction factor SERINC5. J Biol Chem. 2019;

25. Han GZ, Worobey M. An endogenous foamy-like viral element in the coelacanth genome. PLoS Pathog. 2012;

26. Sawyer SL, Wu LI, Emerman M, Malik HS. Positive selection of primate TRIM5 identifies a critical species-specific retroviral restriction domain. Proc Natl Acad Sci. 2005;

27. Newman RM, Hall L, Connole M, Chen G-L, Sato S, Yuste E, et al. Balancing selection and the evolution of functional polymorphism in Old World monkey TRIM5. Proc Natl Acad Sci. 2006;

28. McLaughlin RN, Gable JT, Wittkopp CJ, Emerman M, Malik HS. Conservation and Innovation of APOBEC3A Restriction Functions during Primate Evolution. Mol Biol Evol. 2016;

29. Johnson WE. Rapid adversarial co-evolution of viruses and cellular restriction factors. Curr Top Microbiol Immunol. 2013;

30. Blanco-Melo D, Venkatesh S, Bieniasz PD. Origins and Evolution of tetherin, an Orphan Antiviral Gene. Cell Host Microbe [Internet]. 2016 Aug 10 [cited 2017 Jul 5];20(2):189–201. Available from: http://linkinghub.elsevier.com/retrieve/pii/S193131281630258X

31. Meng X, Zhang F, Yan B, Si C, Honda H, Nagamachi A, et al. A paralogous pair of mammalian host restriction factors form a critical host barrier against poxvirus infection. PLoS Pathog. 2018;

32. Browne EP, Littman DR. Species-Specific Restriction of Apobec3-Mediated Hypermutation. J Virol. 2008;

33. Mitchell PS, Patzina C, Emerman M, Haller O, Malik HS, Kochs G. Evolution-guided identification of antiviral specificity determinants in the broadly acting interferon-induced innate immunity factor MxA. Cell Host Microbe. 2012;

34. Mitchell PS, Young JM, Emerman M, Malik HS. Evolutionary Analyses Suggest a Function of MxB Immunity Proteins Beyond Lentivirus Restriction. PLoS Pathog. 2015;

35. Dutko JA, Schäfer A, Kenny AE, Cullen BR, Curcio MJ. Inhibition of a yeast LTR retrotransposon by human APOBEC3 cytidine deaminases. Curr Biol. 2005;

36. Irwin B, Aye M, Baldi P, Beliakova-Bethell N, Cheng H, Dou Y, et al. Retroviruses and yeast retrotransposons use overlapping sets of host genes. Genome Res. 2005;

37. Miller JE, Zhang L, Jiang H, Li Y, Pugh BF, Reese JC. Genome-wide mapping of decay factor-mRNA interactions in yeast identifies nutrient-responsive transcripts as targets of the deadenylase Ccr4. G3 Genes, Genomes, Genet. 2018;

38. Llorens J V., Clark JB, Martínez-Garay I, Soriano S, De Frutos R, Martínez-Sebastián MJ. Gypsy endogenous retrovirus maintains potential infectivity in several species of Drosophilids. BMC Evol Biol. 2008;

39. Teysset L, Burns JC, Shike H, Sullivan BL, Bucheton A, Terzian C. A Moloney Murine Leukemia Virus-Based Retroviral Vector Pseudotyped by the Insect Retroviral gypsy Envelope Can Infect Drosophila Cells. J Virol. 1998;

40. Mellor J, Fulton SM, Dobson MJ, Wilson W, Kingsman SM, Kingsman AJ. A retrovirus- like strategy for expression of a fusion protein encoded by yeast transposon Ty1. Nature. 1985;

41. Aiewsakun P, Katzourakis A. Marine origin of retroviruses in the early Palaeozoic Era. Nat Commun. 2017;

42. Beitari S, Ding S, Pan Q, Finzi A, Liang C. The effect of HIV-1 Env on SERINC5 antagonism. J Virol [Internet]. 2016;(December):JVI.02214-16. Available from: http://jvi.asm.org/lookup/doi/10.1128/JVI.02214-16

43. Sood C, Marin M, Chande A, Pizzato M, Melikyan GB. SERINC5 protein inhibits HIV-1 fusion pore formation by promoting functional inactivation of envelope glycoproteins. J Biol Chem [Internet]. 2017 Apr 7 [cited 2017 May 26];292(14):6014–26. Available from: http://www.ncbi.nlm.nih.gov/pubmed/28179429

44. Pye, V.E., Rosa, A., Bertelli C et al. A bipartite structural organization defines the SERINC family of HIV-1 restriction factors. Nat Struct Mol Biol. 2020;27:78–83.

45. Beitari S, Ding S, Pan Q, Finzi A, Liang C. Effect of HIV-1 Env on SERINC5 Antagonism. J Virol. 2017;91(4).

46. Sojo V, Dessimoz C, Pomiankowski A, Lane N. Membrane Proteins Are Dramatically Less Conserved than Water-Soluble Proteins across the Tree of Life. Mol Biol Evol. 2016;

47. Li M, Waheed AA, Yu J, Zeng C, Chen HY, Zheng YM, et al. TIM-mediated inhibition of HIV-1 release is antagonized by Nef but potentiated by SERINC proteins. Proc Natl Acad Sci U S A. 2019;

48. Liu G, Zhang H, Zhao C, Zhang H. Evolutionary History of the Toll-Like Receptor Gene Family across Vertebrates. Enard D, editor. Genome Biol Evol [Internet]. 2020 Jan 1 [cited 2020 Feb 7];12(1):3615–34. Available from: https://academic.oup.com/gbe/article/12/1/3615/5652095

49. Simmonds P, Aiewsakun P, Katzourakis A. Prisoners of war — host adaptation and its constraints on virus evolution. Nat Rev Microbiol. 2019;

50. Trobridge G, Josephson N, Vassilopoulos G, Mac J, Russell DW. Improved foamy virus vectors with minimal viral sequences. Mol Ther. 2002;6(3):321–8.

51. Pizzato M, Erlwein O, Bonsall D, Kaye S, Muir D, McClure MO. A one-step SYBR Green I-based product-enhanced reverse transcriptase assay for the quantitation of retroviruses in cell culture supernatants. J Virol Methods [Internet]. 2009 Mar [cited 2017 Jun 19];156(1–2):1–7. Available from: http://www.ncbi.nlm.nih.gov/pubmed/19022294

52. Mittal P, Jaiswal SK, Vijay N, Saxena R, Sharma VK. Comparative analysis of corrected tiger genome provides clues to its neuronal evolution. Sci Rep [Internet]. 2019 Dec 5 [cited 2020 Jan 13];9(1):18459. Available from: http://www.nature.com/articles/s41598-019-54838-z

53. Li H. [Heng Li - Compares BWA to other long read aligners like CUSHAW2] Aligning sequence reads, clone sequences and assembly contigs with BWA-MEM. arXiv Prepr arXiv. 2013;

54. Korneliussen TS, Albrechtsen A, Nielsen R. ANGSD: Analysis of Next Generation Sequencing Data. BMC Bioinformatics. 2014;

55. Thomas GWC, Hahn MW. Referee: Reference assembly quality scores. Genome Biol Evol. 2019;

56. Shinde SS, Teekas L, Sharma S, Vijay N. Signatures of Relaxed Selection in the CYP8B1 Gene of Birds and Mammals. J Mol Evol. 2019;

57. Blackburne BP, Whelan S. Class of multiple sequence alignment algorithm affects genomic analysis. Mol Biol Evol. 2013;

58. Murrell B, Moola S, Mabona A, Weighill T, Sheward D, Kosakovsky Pond SL, et al. FUBAR: A fast, unconstrained bayesian AppRoximation for inferring selection. Mol Biol Evol. 2013;

59. Kosakovsky Pond SL, Frost SDW, Muse S V. HyPhy: Hypothesis testing using phylogenies. Bioinformatics. 2005;

60. Avino M, Poon AFY. Detecting Amino Acid Coevolution with Bayesian Graphical Models. In: Methods in Molecular Biology. 2019.

