## Supplementary material for "Coevolution of retroviruses with *SERINC*s following whole-genome duplication divergence": SI FIgures 1-15

**Fig-S1**. **Phylogenetic analysis of SERINC paralogs**. *S. cerevisiae, C. elegans,* and *D. melanogaster* has a single copy of this gene. Whole-genome duplication that occurred in an early ancestor of mammals gave rise to a cluster of five genes divided into clusters of SERINC1, SERINC2, SERINC3, and SERINC4 and SERINC5. Colors denote different branches of each SERINC in chordates. The lengths of the triangles are proportional to the number of nucleotide substitutions that have taken place in a particular branch. The tree was generated using Ensembl.

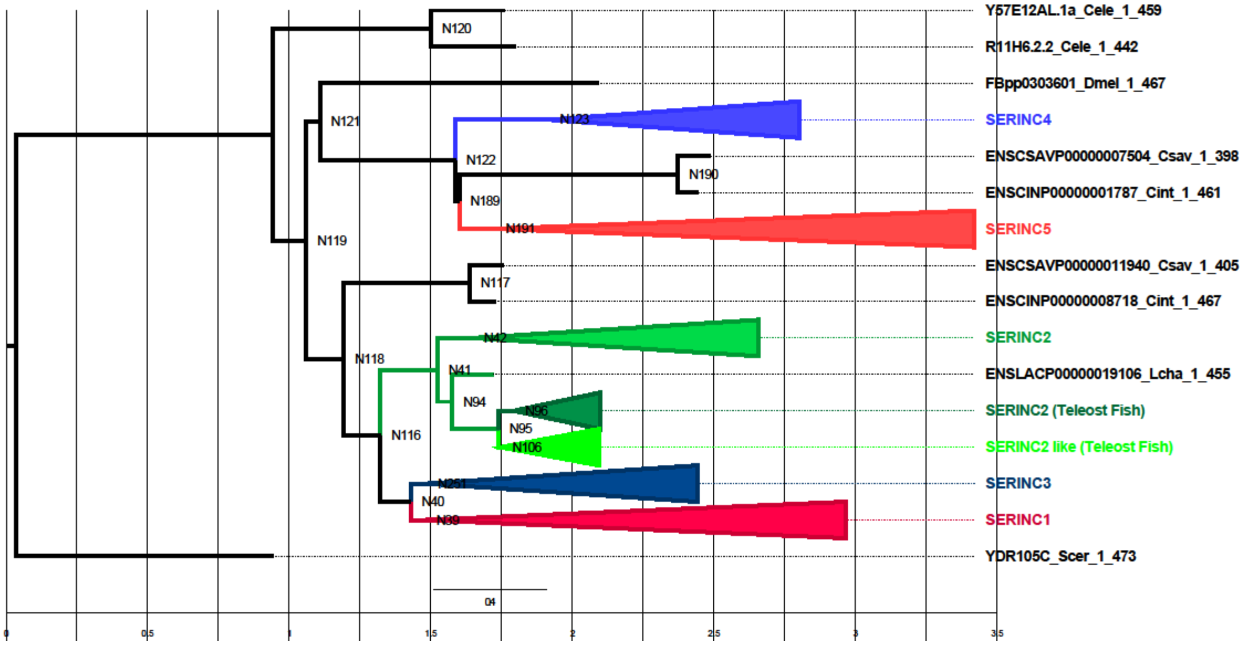

**Fig-S2**. Topology of human SERINC Paralogs predicted using TOPCONS

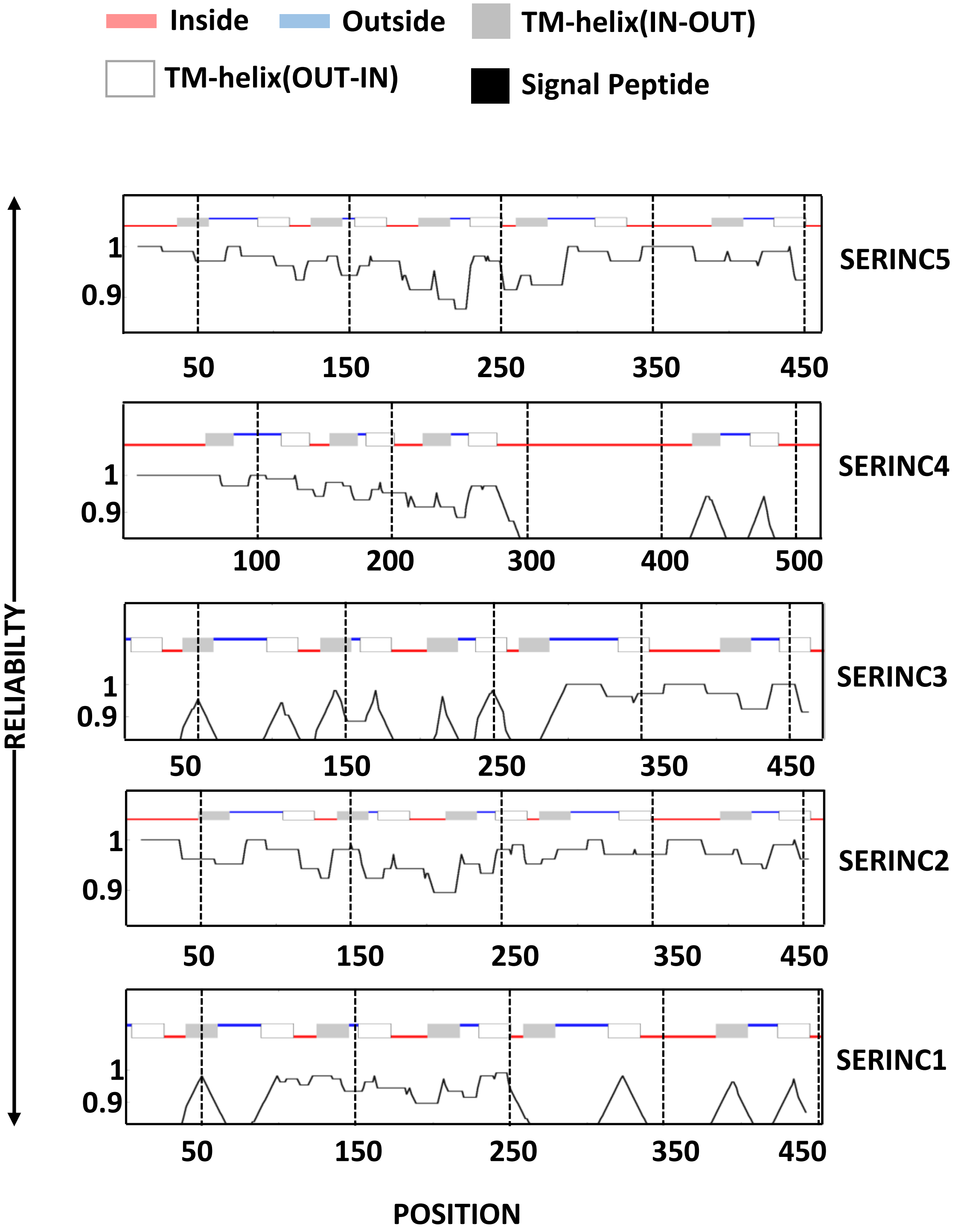

**Fig-S3**. Arms-race signatures of primate SERINC genes inferred using FUBAR

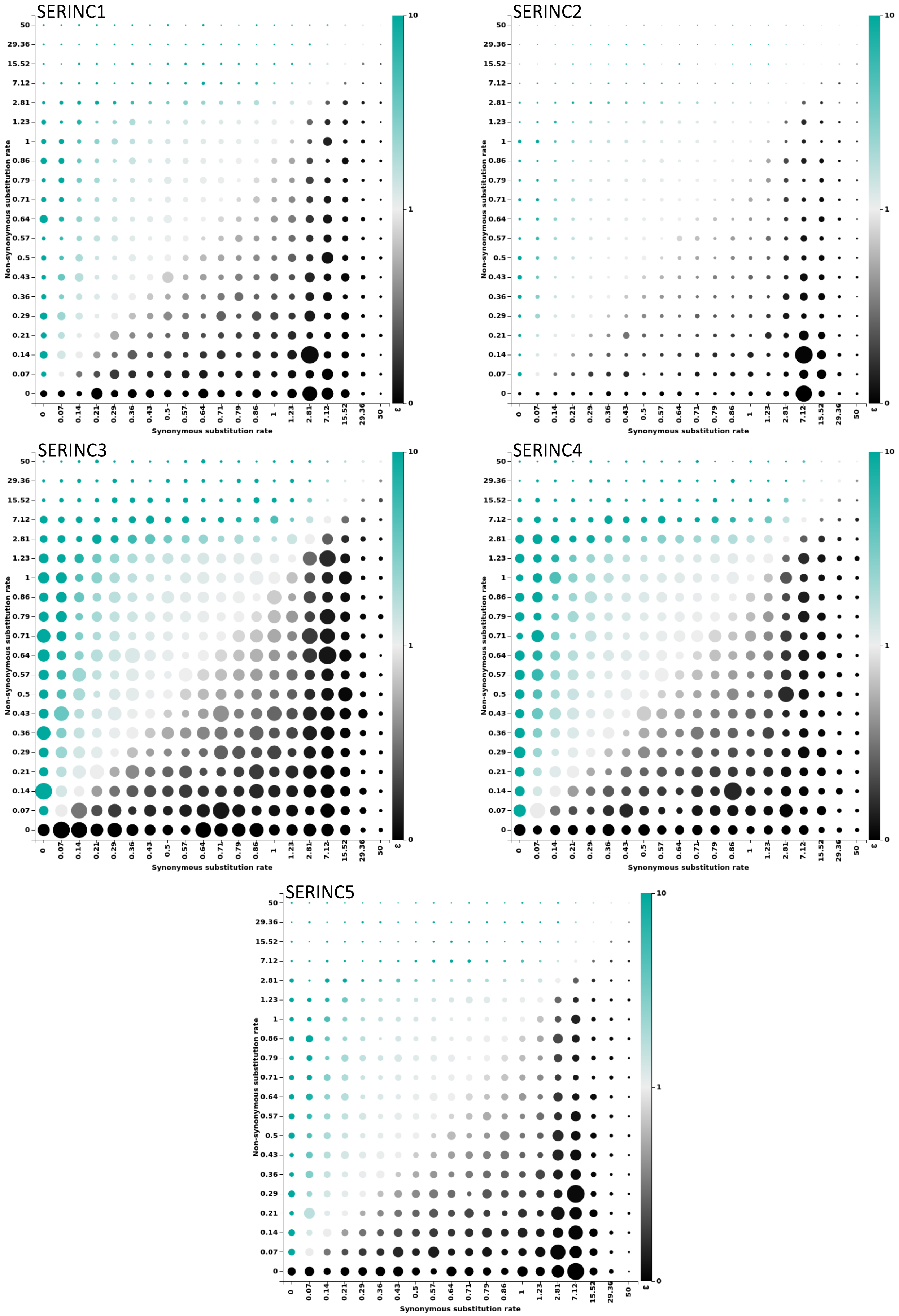

**Fig-S4**. Distribution of sites under different selection regimes across the SERINC genes.
The colors are inner (Blue), outer (Pink), helix (Red). Black dots are estimates of dS and red dots are dN. Pairs of conditionally dependent sites identified by the program BGM (Bayesian Graphical Model) are connected by a dotted line (see SERINC1 and SERINC4). Vertical green lines correspond to the sites under purifying selection.

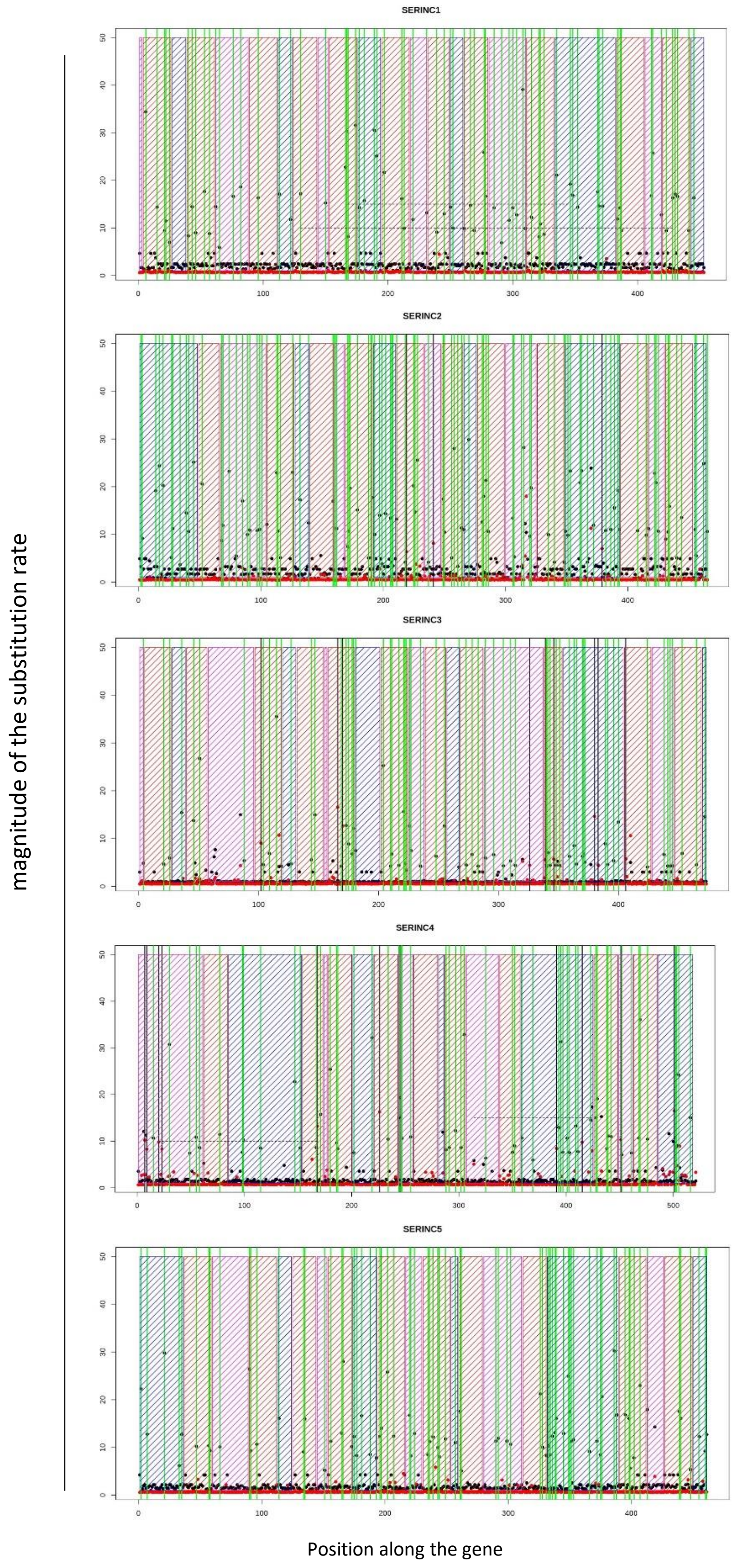

**Fig-S5**. Effect of different aligners on hierarchical clustering of arms-race signatures of primate genes. Interferon induced genes identified by (Shaw et al 2017) are color coded as red (upregulated), blue(downregulated), green (SERINC paralogs), grey (other select genes)

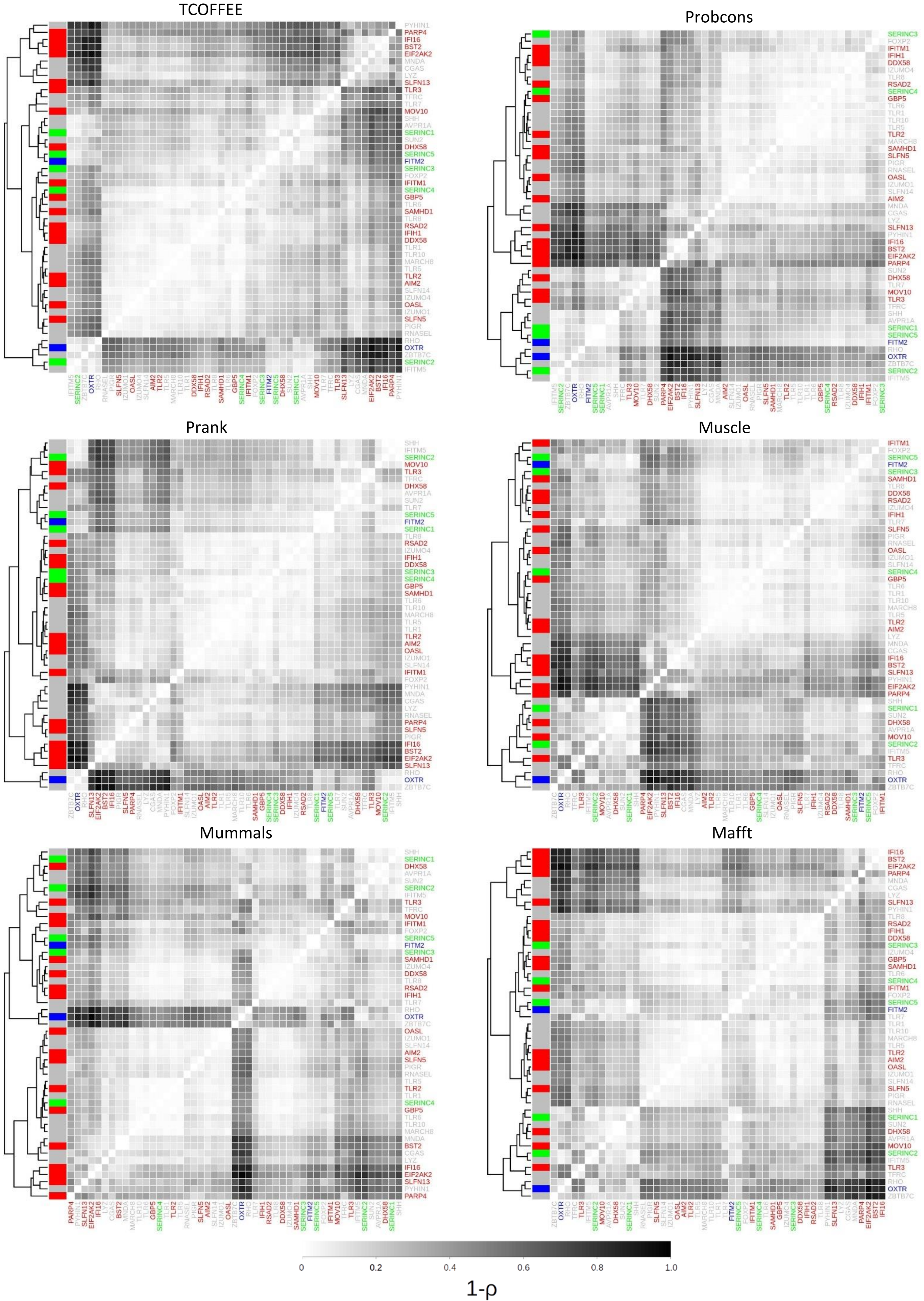

**Fig-S6.** Topology of SERINC5 orthologs predicted using Topcons. Coelacanth SERINC5 was reconstructed from the RNAseq data (*) see supplemental material for details

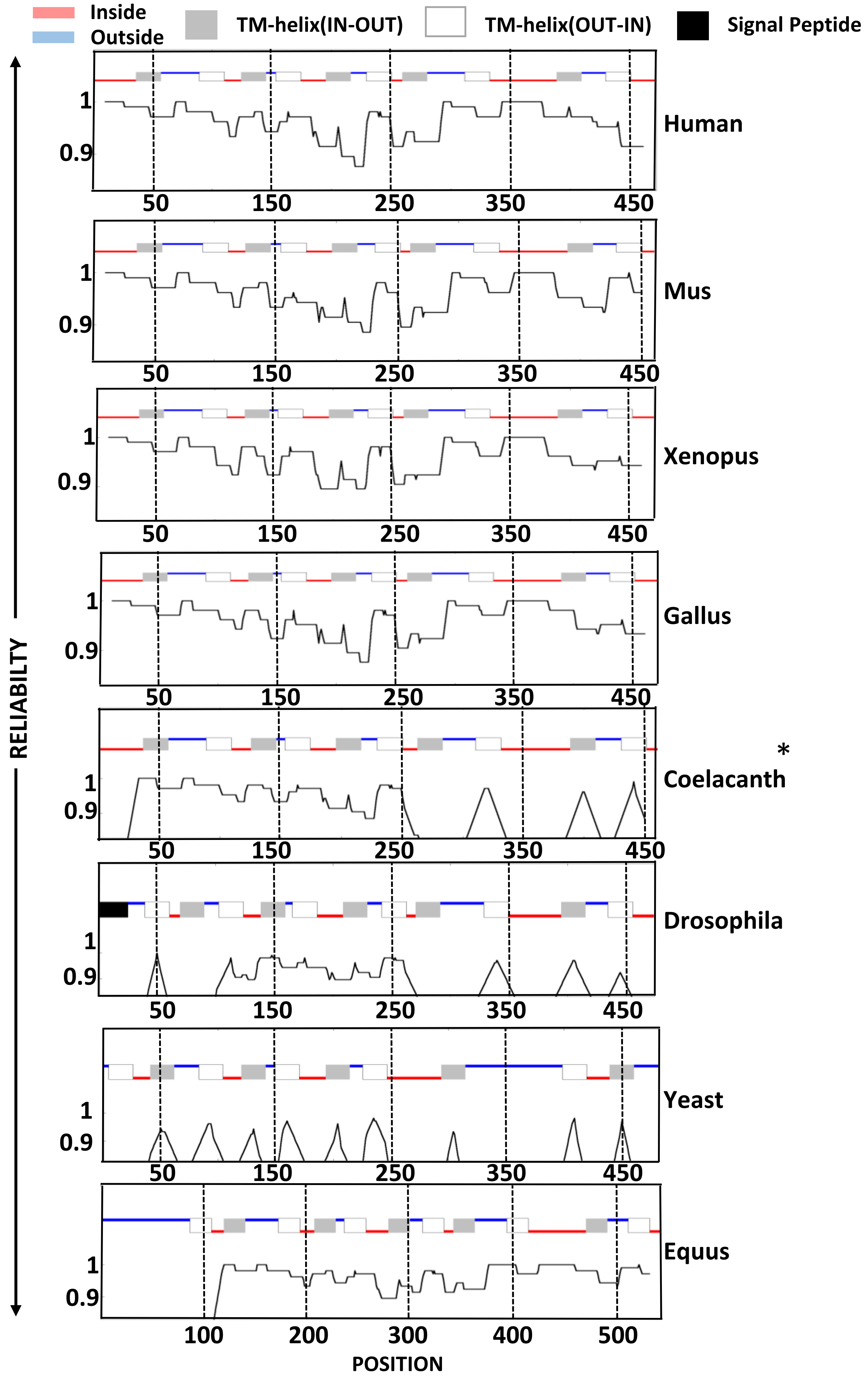

**Fig-S7**. Topological features of Human SERINC2 splice isoforms and coelacanth SERINC2 predicted using TOPCONS

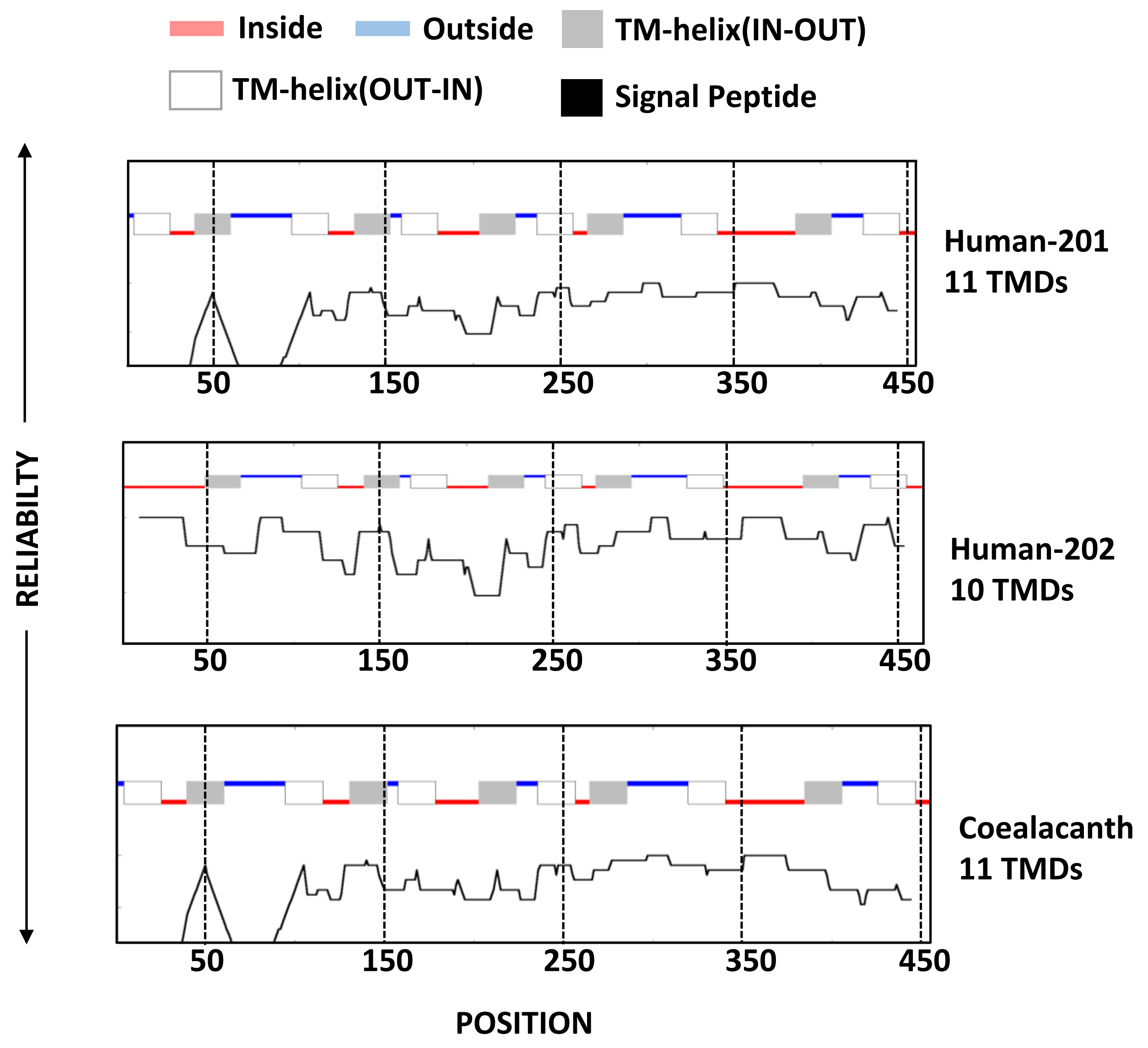

**Fig-S8**. Topology of SERINC2 orthologs predicted using TOPCONS

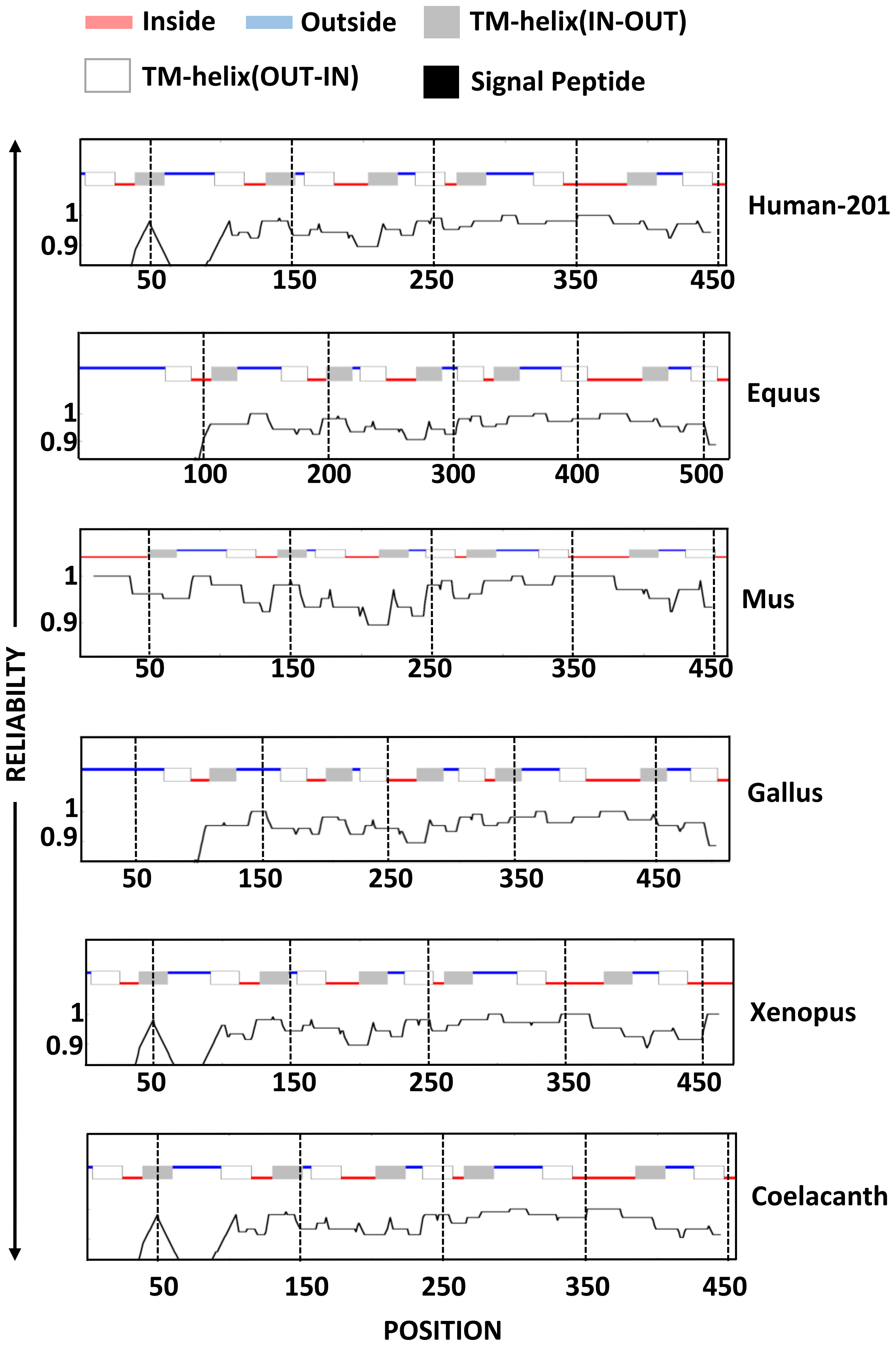

**Fig-S9**. Incorporation of SERINC2 orthologs in the virus particles and detection by indicted antibodies against target proteins

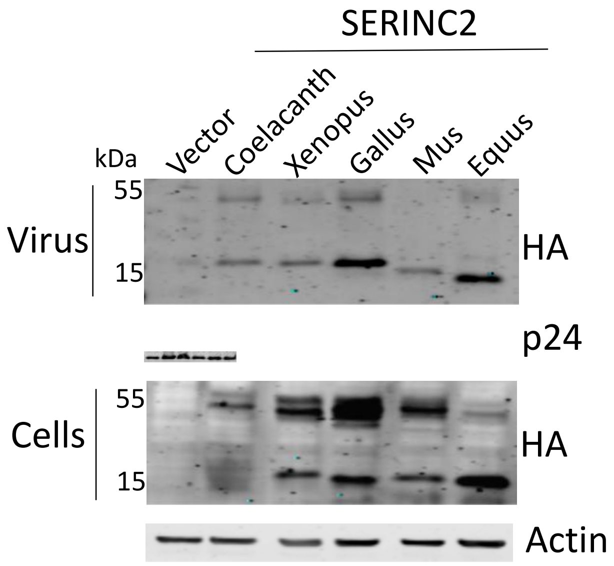

**Fig-S10**. Effect of a dose-dependent expression of coelacanth SERINC2 on HIV-1 infectivity

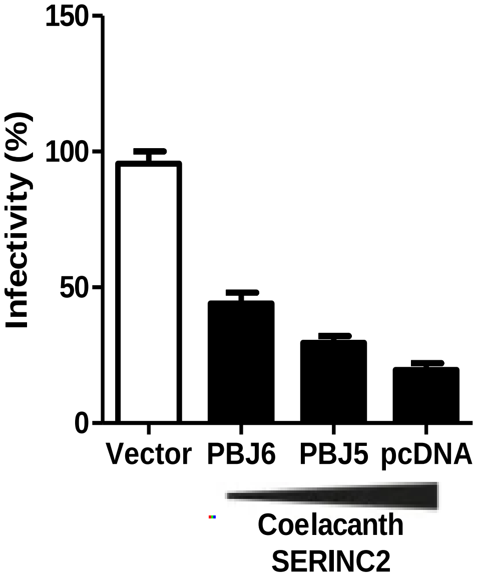

**Fig-S11**. Counteraction of coelacanth SERINC2 restriction by a Foamy virus envelope glycoprotein

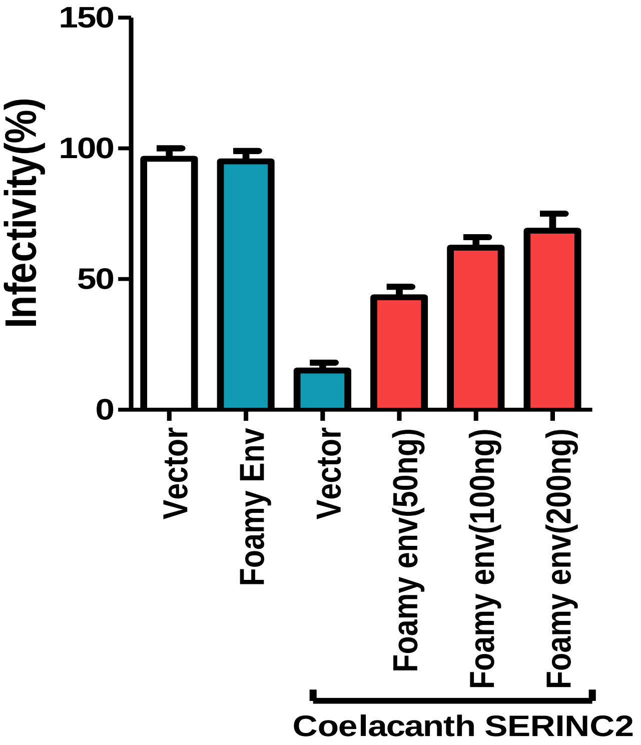

**Fig-S12**. Reciprocal-packaging of a foamy envelope with an HIV core and sensitivity to SERINCs

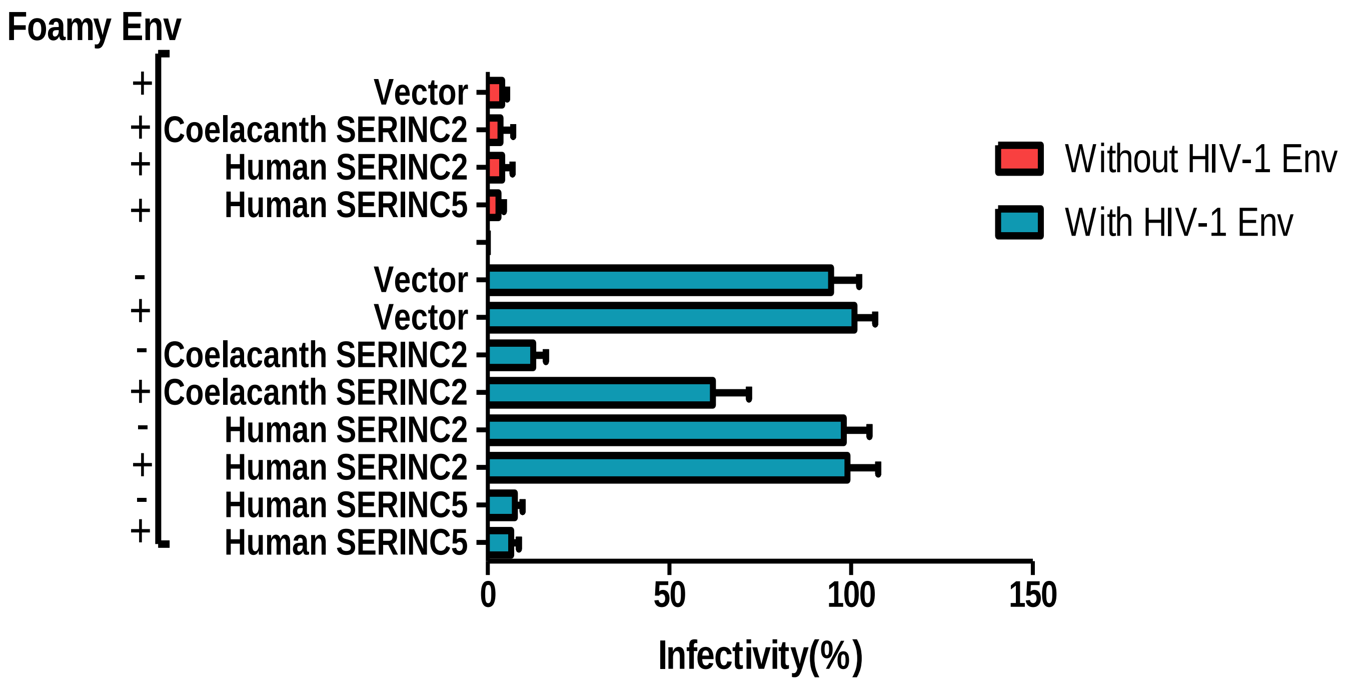

**Fig-S13**. Infectivity of HIV-1 produced from JTAg$^{SERINC5/3KO}$ having ectopic expression of the indicated SERINCs

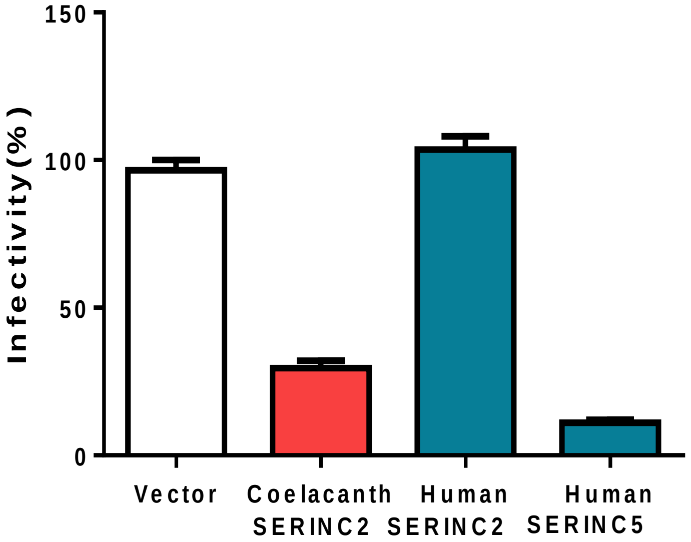

**Fig-S14**. Multiple sequence alignment of SERINC2 orthologs; highlighted regions showing the sites of sequence divergence

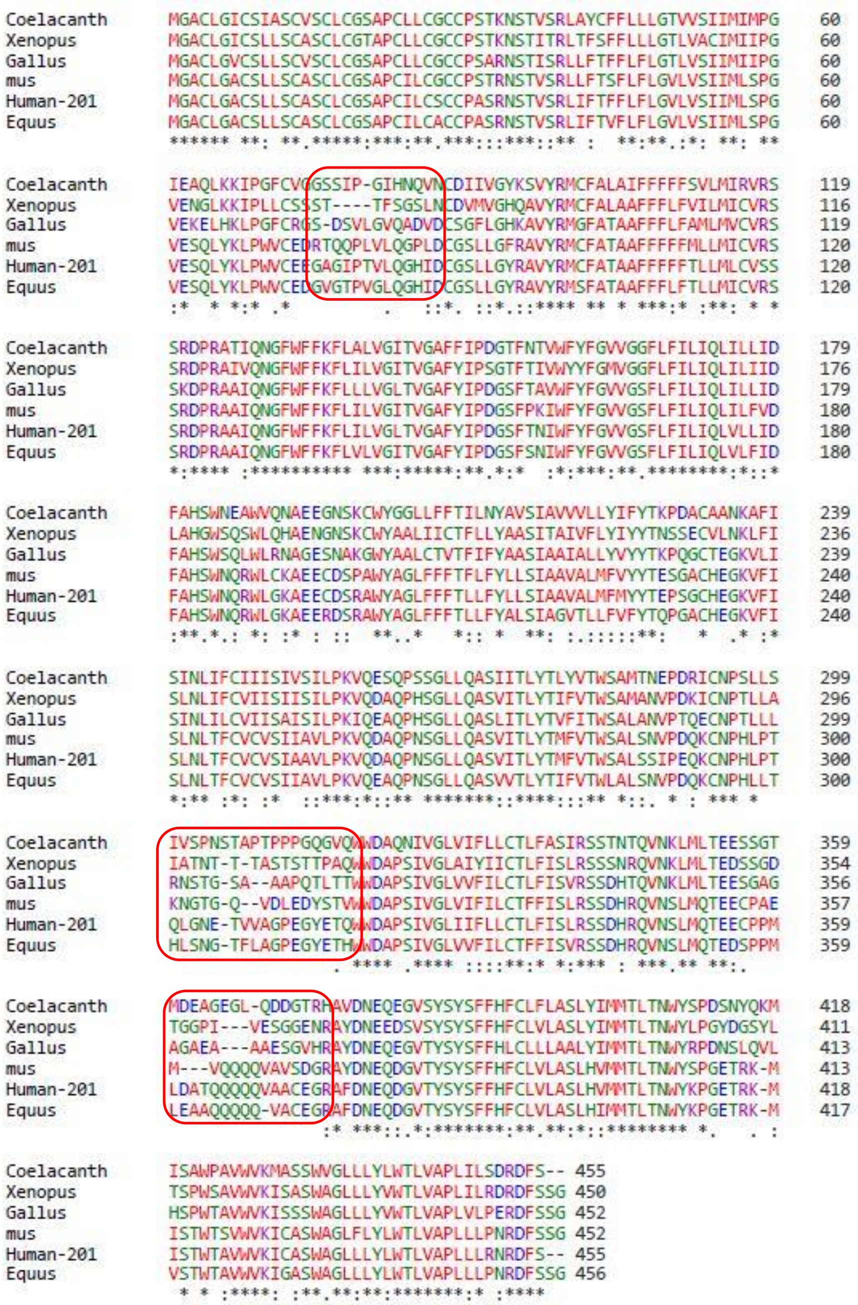

**Fig-S15**. Evidence for the presence of HNF4a binding site in the intron of human SERINC2 gene

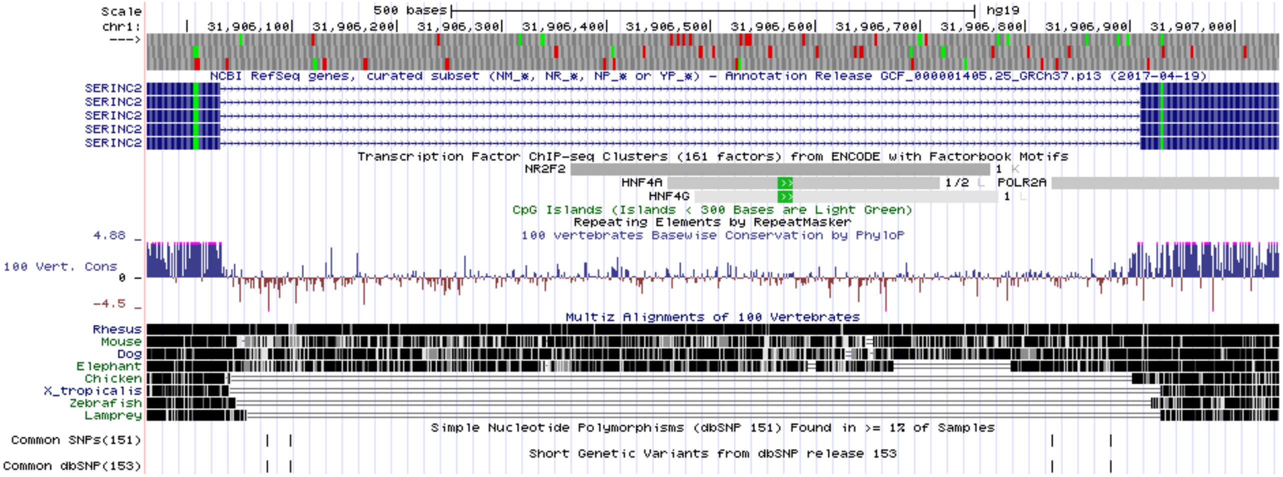
