## Supplementary figures and images for "Coevolution of retroviruses with *SERINC*s following whole-genome duplication divergence"

### SI Figure 16

**Fig-S16.** Gene expression evolution of SERINC genes in vertebrates

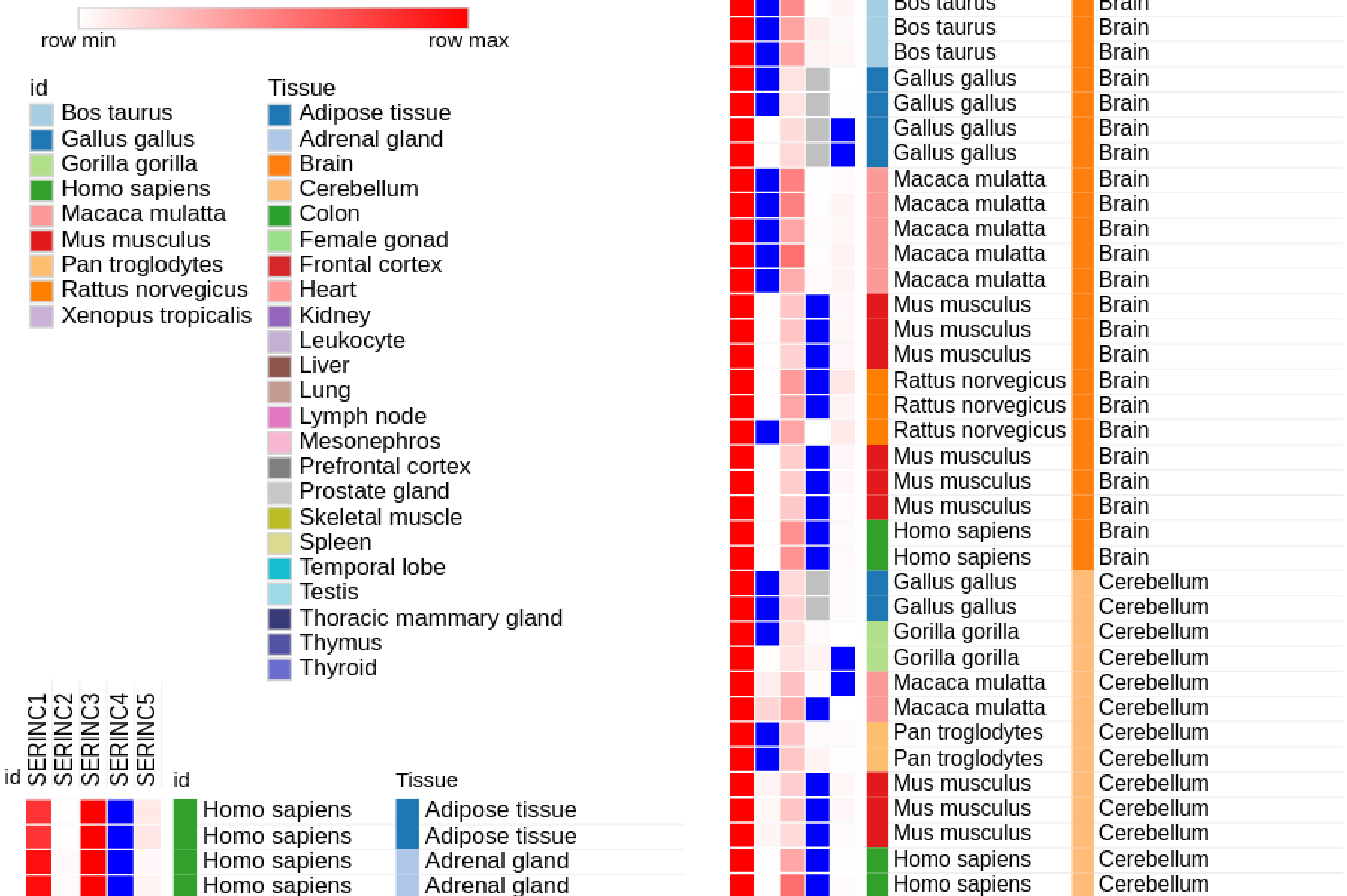

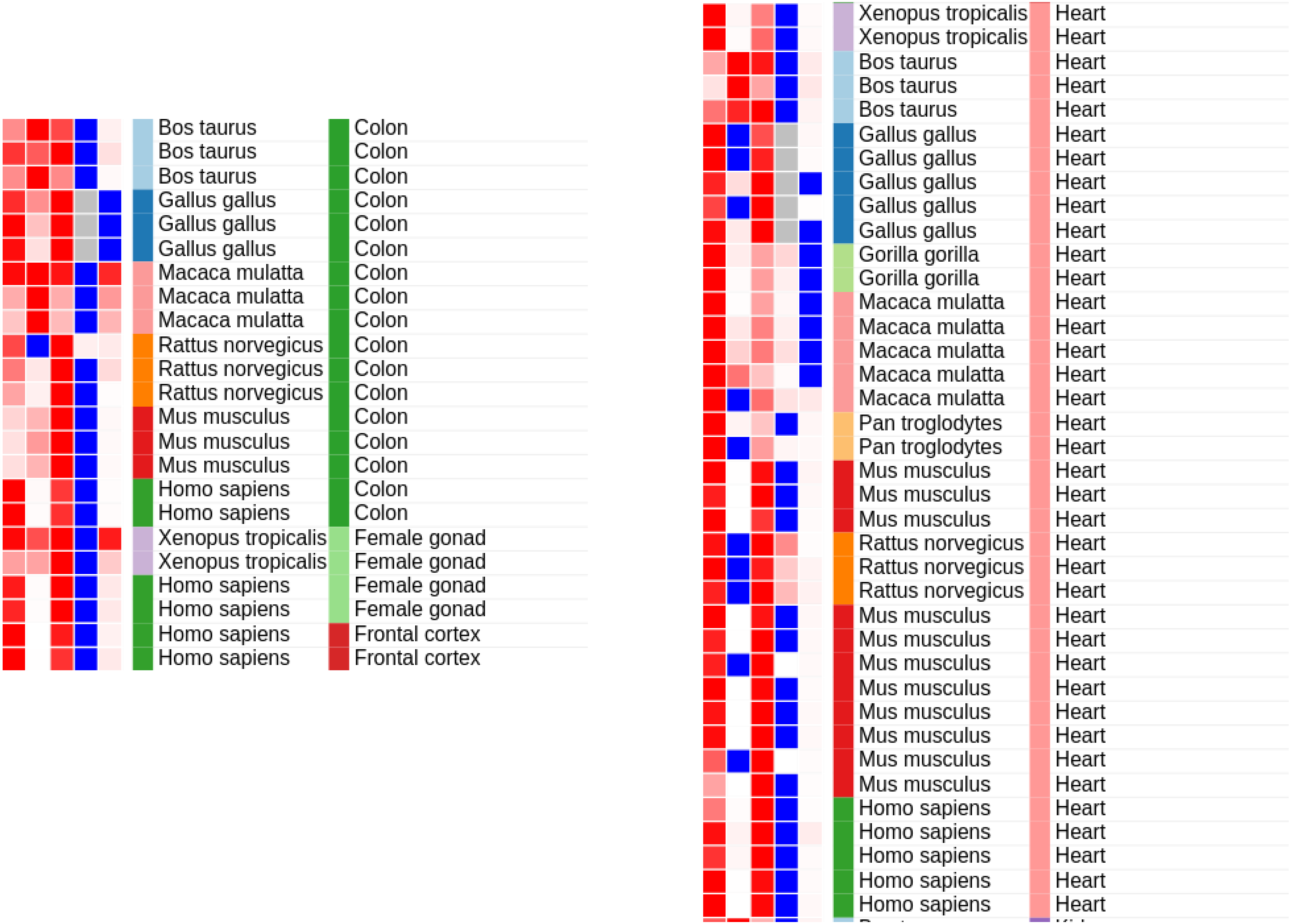

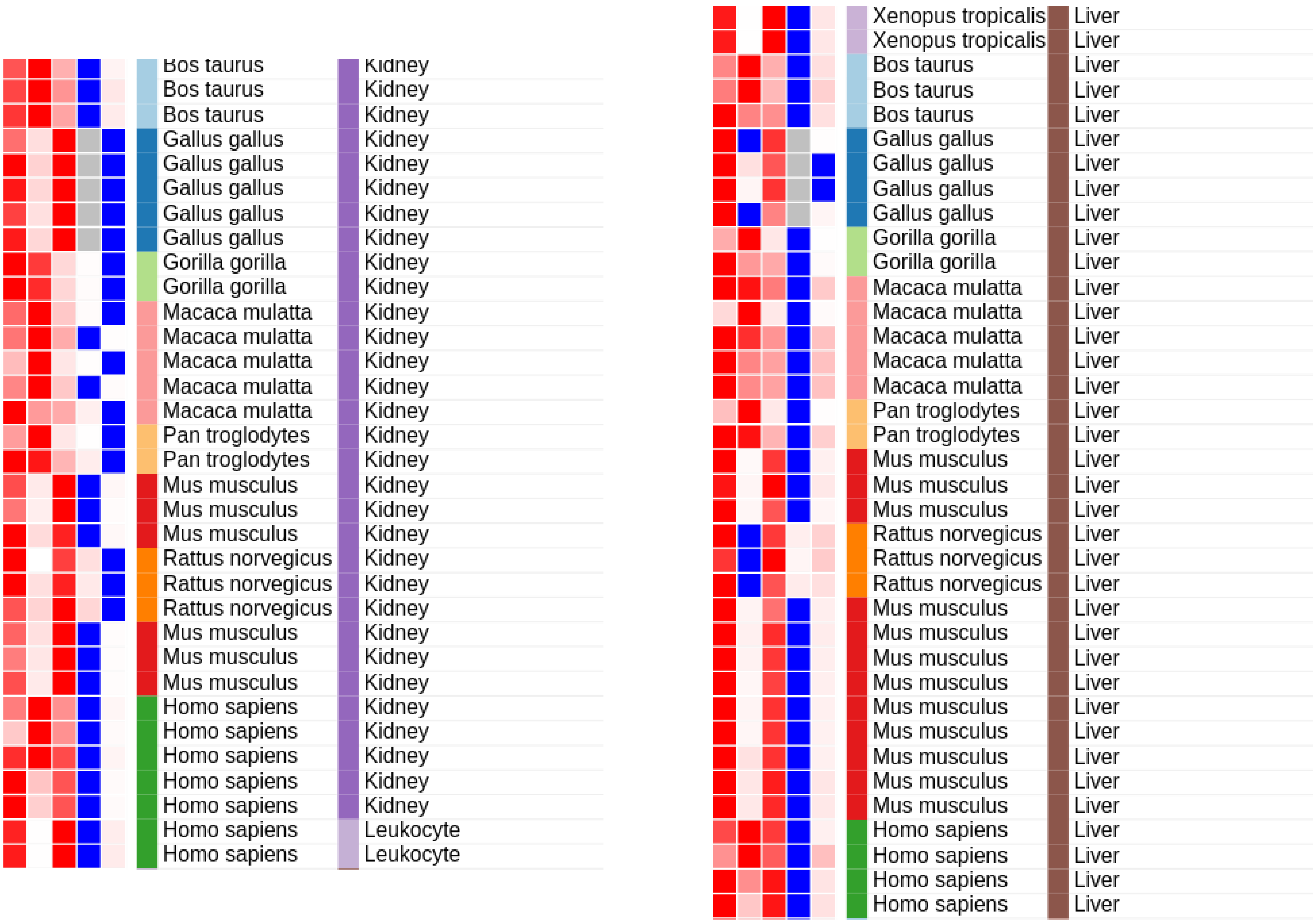

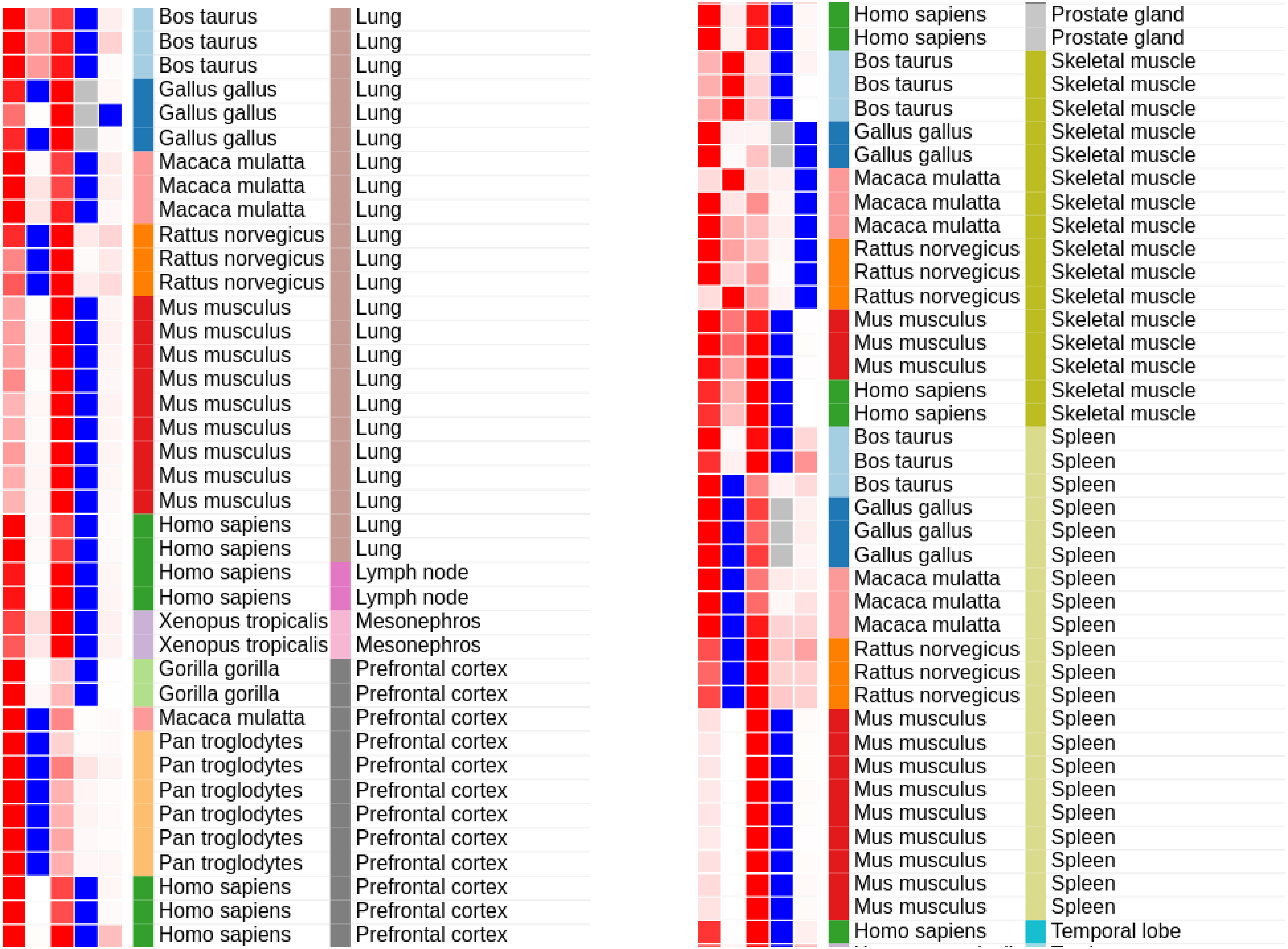

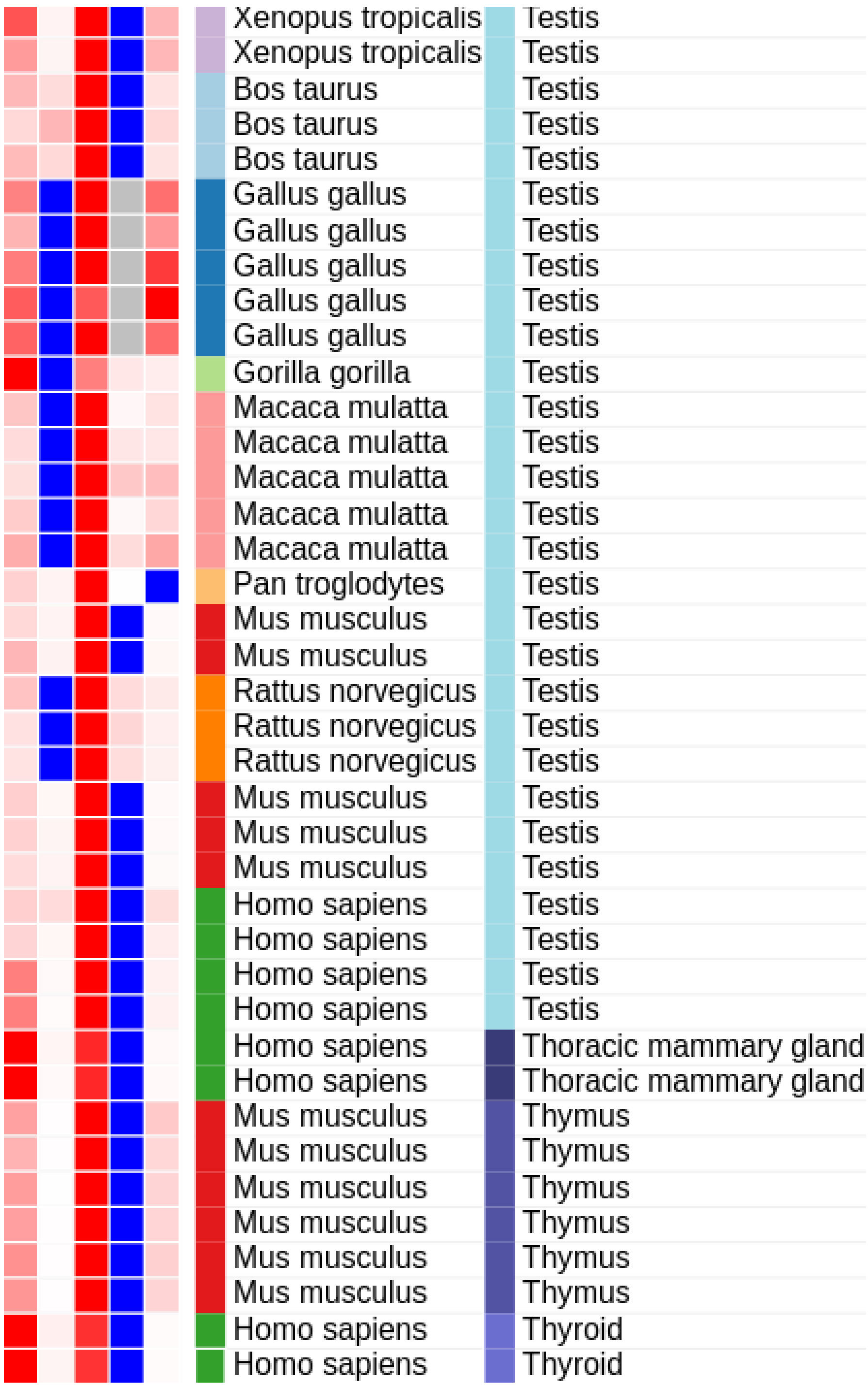
